# Gene expression variability in human and chimpanzee populations share common determinants

**DOI:** 10.1101/2020.06.11.146142

**Authors:** Benjamin J. Fair, Lauren E. Blake, Abhishek Sarkar, Bryan J. Pavlovic, Claudia Cuevas, Yoav Gilad

## Abstract

Inter-individual variation in gene expression has been shown to be heritable and it is often associated with differences in disease susceptibility between individuals. Many studies focused on mapping associations between genetic and gene regulatory variation, yet much less attention has been paid to the evolutionary processes that shape the observed differences in gene regulation between individuals in humans or any other primate. To begin addressing this gap, we performed a comparative analysis of gene expression variability and expression quantitative trait loci (eQTLs) in humans and chimpanzees, using gene expression data from primary heart samples. We found that expression variability in both species is often determined by non-genetic sources, such as cell-type heterogeneity. However, we also provide evidence that inter-individual variation in gene regulation can be genetically controlled, and that the degree of such variability is generally conserved in humans and chimpanzees. In particular, we found a significant overlap of orthologous genes associated with eQTLs in both species. We conclude that gene expression variability in humans and chimpanzees often evolves under similar evolutionary pressures.

## Introduction

Variation in gene expression underlies many phenotypic differences between and within species. Gene expression itself is a quantitative phenotype that is subject to both random drift and natural selection. A deeper understanding of how natural selection shapes gene expression across primates is central to our understanding of human evolution, and may also elucidate the mechanistic basis for variation in quantitative traits and disease risk within species.

Several studies have used a comparative transcriptomics approach across species and tissues to identify genes whose expression patterns are consistent with the action of natural selection (Barbosa-Morais et al., 2012; Brawand et al., 2011; Merkin et al., 2012). For example, a pattern of highly similar gene expression levels in all primates may be consistent with the action of stabilizing selection on gene regulation, and potentially, negative selection on the corresponding regulatory elements. Indeed, many studies have found that the expression of most genes evolves slower than expected under neutrality (Chan et al., 2009; Khaitovich et al., 2006; Khan et al., 2013; Lemos et al., 2005; Merkin et al., 2012; Romero et al., 2012). Conversely, genes that show a reduced or elevated expression level exclusively in the human lineage may indicate directional selection on gene regulation in humans, potentially resulting in positive selection for particular regulatory variants (Blekhman et al., 2008; Gilad et al., 2006; Perry et al., 2012). However, it is often difficult to determine whether lineage-specific expression changes are due to inter-species environmental differences or to natural selection on certain regulatory variants.

A complementary approach to understanding gene regulation and associated selection pressures utilizes within-species variation to identify genetic variants that affect gene expression levels. Such variants are referred to as expression quantitative trait loci, or eQTLs. Overall, there is some evidence that eQTLs are evolving under weak negative selection. For example, the magnitude of eQTL effect size on expression levels is weakly anti-correlated with eQTL allele frequency (Battle et al., 2014; Glassberg et al., 2019). Additionally, the set of human genes associated with an eQTL (eGenes) tend to be slightly depleted for genes relevant to universally conserved and essential cellular processes (Popadin et al., 2014; Tung et al., 2015; Ward and Gilad, 2019), and for genes at the center of protein interaction networks (Battle et al., 2014; Mähler et al., 2017). These observations are consistent with negative selection purging strong regulatory variants within species, particularly variants that regulate the expression of genes whose precise regulation is essential.

The eQTL mapping approach allows us to connect genetic variants to the genes they regulate, and provides a mechanistic explanation for a portion of the heritability of gene expression phenotypes (Price et al., 2011; Wright et al., 2014). However, not all gene expression level variation can be explained by genetic variation, and of the genetic contribution, only about 20% of heritability can be explained by locally-acting variants (referred to as *cis* eQTLs; (Albert et al., 2018; Price et al., 2011). We assume that the remaining 80% of heritable expression variation is determined by distally-acting regulatory variants (*trans* eQTLs). However, because *trans* eQTLs can be located anywhere in the genome, they are difficult to pinpoint, even in studies with large sample sizes (Battle et al., 2014; Westra et al., 2013). Therefore, regarding the identification of genes undergoing stabilizing selection within or between closely related species, the insights gained by *cis* eQTL approaches may be limited.

A third approach to understanding the evolutionary forces on gene expression is to directly quantify gene expression variability (as opposed to the mean value) within or between populations or species. The variability of gene expression in a population is the sum of variation introduced by local and distal genetic regulation, as well as other sources, including epigenetic (Bashkeel et al., 2019) and environmental effects. The variation in gene expression observed in a given population is the result of all of these effects as well as any technical variation that was introduced during experimental data collection. While variability measurements alone cannot help disentangle the genetic component of variation from the non-genetic component, we do expect that natural selection will minimize the regulatory variation of genes with dosage-sensitive functions. Fortunately, it is possible to obtain relatively stable measurements of population variability for every expressed gene using a moderate sample size of just tens of individuals (de Jong et al., 2019).

The quantification of regulatory variability may provide a window into distinct biological phenomena that are difficult to ascertain by studying inter-species differences in mean expression levels. For example, the adaptability of a population in response to new stresses may be a general function of gene expression variability, as has been demonstrated in yeast populations (Bódi et al., 2017; Wang and Zhang, 2011; Zhang et al., 2009). Furthermore, identification of hypervariable genes, regardless of the genetic or non-genetic source of variability, may help identify genes and pathways which confer differences in disease susceptibilities and treatment responses (Ho et al., 2008; Knowles et al., 2018; Simonovsky et al., 2019). Direct measurements of variability may therefore be a useful complement to the analysis of differences in mean gene expression levels or to eQTL mapping, and could contribute to a better understanding of gene expression evolution and complex traits.

A previous analysis of gene expression variability within and between human populations found higher regulatory variation in genes associated with disease susceptibility (Li et al., 2010). This study did not otherwise identify any particular functional classes or features of genes that showed significant inter-population differences in expression variability – although given the relative genetic similarity, short evolutionary timescale, and migrations between human populations, we might not expect different signatures of selection to be apparent. Nonetheless, this study found that regulatory variability across genes correlates with the levels of genetic variability at nearby loci (Li et al., 2010), thereby providing some measure of support to the notion that differences in regulatory variability between genes is at least partly genetically encoded.

We sought to better understand selection pressures on gene regulation variability by collecting gene expression data from humans and chimpanzees (*Pan troglodytes*). The availability of suitable samples from chimpanzees has notoriously been a limitation for comparative functional genomic studies. Here, we were able to collect the largest population sample of chimpanzee primary tissues to date, allowing not only for sensitive assessment of differences in mean expression levels between species, but also differences in variability within species. To better isolate the genetic component of variability, we complemented the analysis of gene expression levels with a comparative eQTL mapping approach.

## Results

We analyzed RNA-seq data from post-mortem primary heart tissue samples of 39 human and 39 chimpanzee individuals (TableS1). Data from 11 of these humans and 18 of the chimpanzees were previously collected in our lab and published (Pavlovic et al., 2018). We obtained additional human data from the GTEx consortium (post-mortem heart left ventricle), filtering for high quality samples with sufficient read depth (FigS1A-B, see Methods for filtering criteria). To complement the published data from chimpanzees, we generated new RNA-seq data from primary heart samples of 21 additional chimpanzee individuals (see Methods). Because the data were collected at different times, and some of the data from humans were collected in different labs (the GTEx data), our overall study design introduces clear technical batch effects that are partly confounded with species (FigS1B). Our focus, however, is on inter-species comparisons of gene expression variability and eQTLs, which are estimated and identified using the within-species data. The batch effects are therefore expected to have only a minimal impact on the reported results.

To analyze the RNA-seq data, we employed a uniform processing pipeline that only considers reads mapping to human-chimpanzee orthologous exonic regions (Pavlovic et al., 2018). Using these mapped reads, we estimated gene expression levels for each individual in each species. We excluded lowly expressed genes (see Methods), and retained data from 13,432 expressed genes for further analysis. Principle component analysis (FigS1C) and unsupervised clustering (FigS1B) of the data show that, as expected, samples primarily separate by species, and not by source or batch, though as we pointed out, some technical batches are partly confounded with species in this study.

**FigS1.**
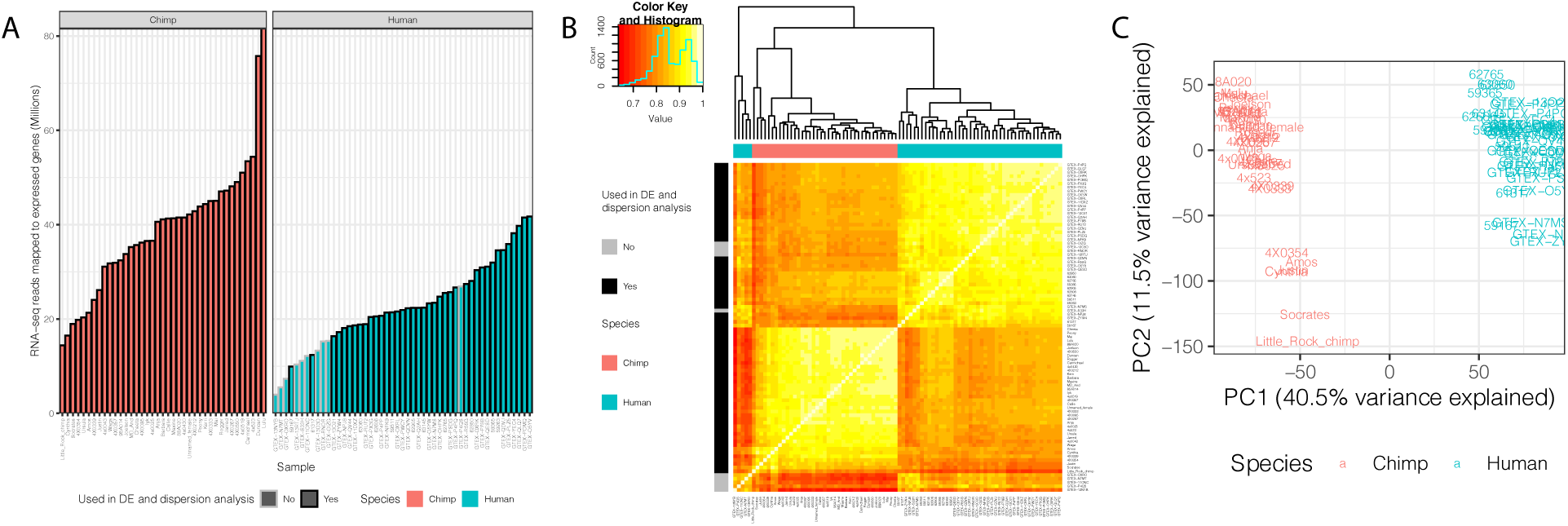
Summary of RNA-seq datasets. (A) Number of reads mapped to expressed genes for each sample. To obtain a balanced set of 39 samples per species, the 10 samples outlined in gray were excluded from differential expression and variability analysis. Nine of these samples are among the lowest read count samples sourced from GTEx. The remaining sample was excluded on the basis of inspection of (B) unsupervised hierarchal clustering of RNA-seq samples by Pearson correlation matrix of expression (CPM). (C) Principal component analysis shows samples separating by species along the first principal component. Only samples used in DE and dispersion analyses are shown.

Using a linear model framework (see Methods), we identified 8,880 differentially expressed (DE) genes between species (FDR<0.05; TableS2), including 6,409 DE genes with effect sizes smaller than a twofold change (FigS2). As our comparative sample size is unusually large, we have an opportunity to comment on the robustness of observations that are made using study designs with smaller sample sizes, which are more typical for comparative studies in primates (Khan et al., 2013; Pavlovic et al., 2018; Perry et al., 2012). To do so, we subsampled our data and repeated the DE analysis for various sample sizes and read depths. To benchmark the results, we used the effect size estimates and DE gene classifications of the full dataset as an *ad hoc* gold standard reference.

**FigS2.**
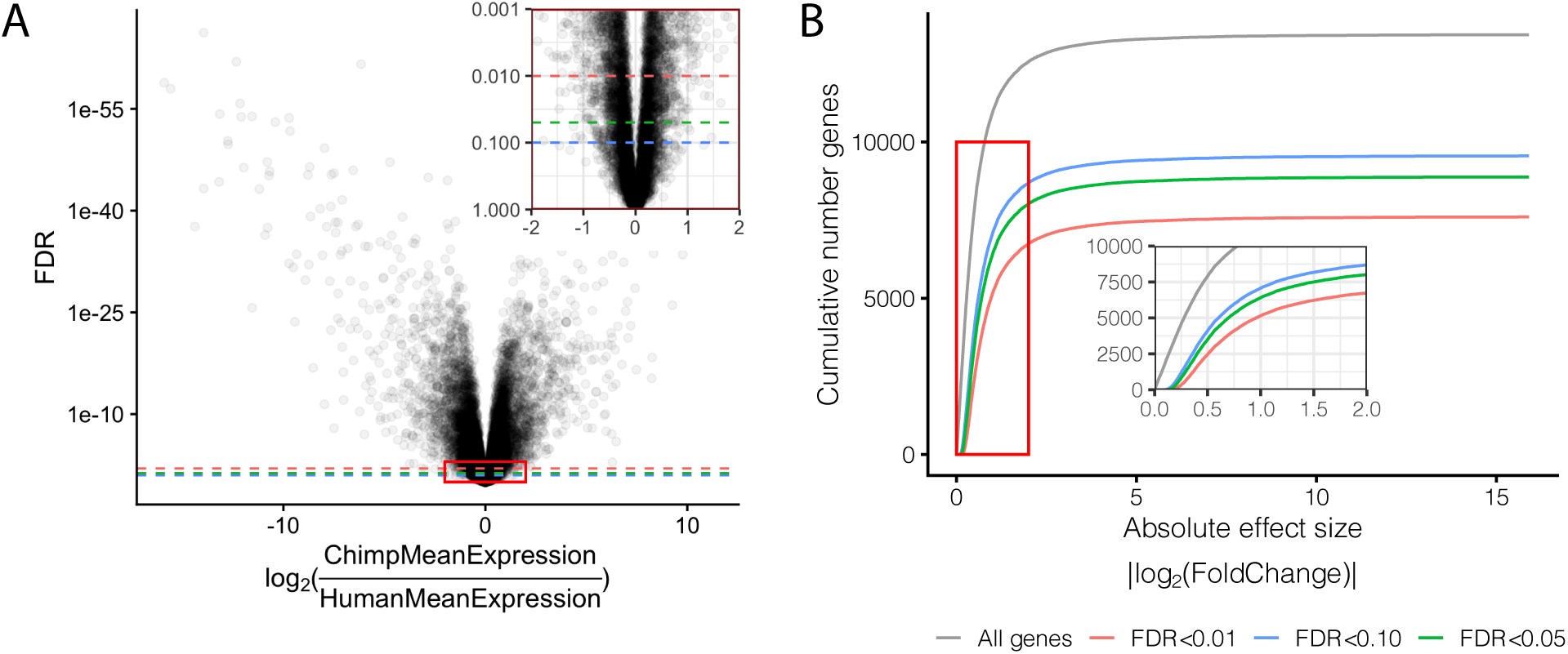
Effect size and significance of differential expression (DE) analysis. (A) Volcano plot of DE genes between human and chimpanzee heart tissue. Inset focuses on the red boxed area to highlight the relationship between expression fold change and DE gene classification at various FDR thresholds. (B) The number of DE genes identified under any given fold change threshold at various FDR thresholds.

As expected, we found that the number of individuals, not sequencing depth (within the range of 10M to >25M reads per sample), is the primary driver of power to detect inter-species differences in gene expression (FigS3F-O). Our results indicate that while comparative studies with small sample sizes are underpowered, their reports of differentially expressed genes are generally robust. For example, when we used a sample size of only 4 chimpanzee individuals and 4 human individuals, we identified a median of 1,373 (869 – 2,131 interquartile range across 100 resamples) DE genes (FDR<0.05), or just 15% (10 – 24% IQR) of the genes identified as DE in the analysis of the full set of samples (FigS3A). Furthermore, a study with only 4 individuals from each species is particularly underpowered to identify subtle inter-species differences (FigS3D), only capturing a median of 5% (2 – 12% IQR) of the DE genes with effect sizes smaller than a twofold change. In addition, of the DE genes identified at this sample size, the estimated magnitude effect sizes are typically upwardly biased (FigS3E) due to the winner’s curse effects (Göring et al., 2001; Ioannidis, 2008), which particularly affect under-powered study designs (FigS3E). However, even at this small sample size, the false positive rate associated with the classification of DE genes is well calibrated; when we used an FDR of 5% to classify genes as DE between the species, we empirically estimated a median of 2% (1 – 5% IQR) false discoveries based on the gold standard reference. Similar analyses for study designs with different sample sizes are available in FigS3.

**FigS3.**
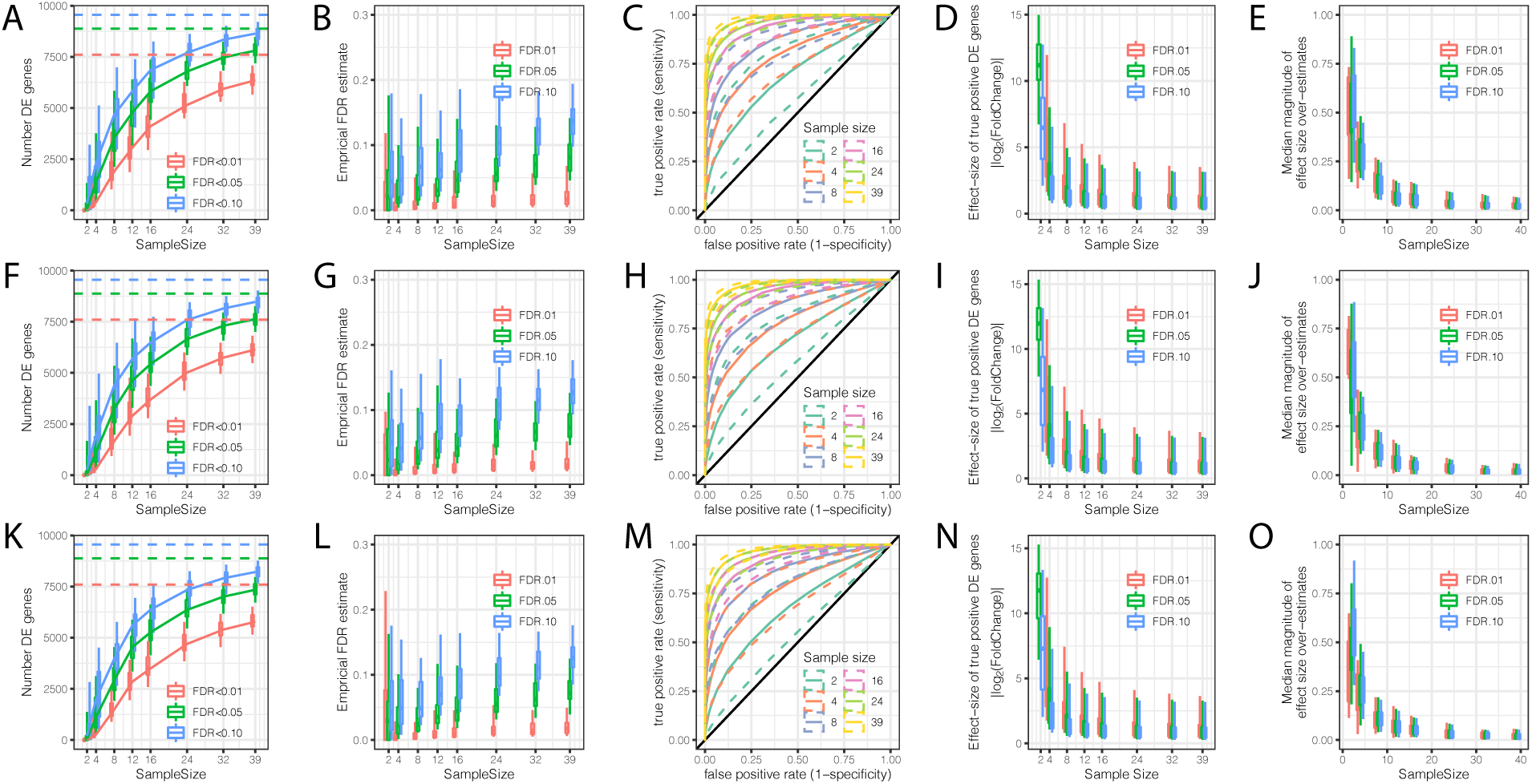
Evaluation of the contribution of sample size and read depth to differential expression analysis between chimpanzee and human. (A) The number of DE genes identified at varying thresholds after randomly subsampling (with replacement) the number of individuals in each species at the indicated sample size per species (hereafter referred to as a single bootstrap replicate). Dashed lines indicate the number of DE genes identified from the full dataset of 39 human and 39 chimpanzee individuals. Box-whisker plots depict the 0.05, 0.25, 0.5, 0.75, and 0.95 quantiles among 100 bootstrap replicates. (B) An empirical estimate of FDR was obtained by calculating the fraction of DE genes in each subsample which were not identified at FDR<0.01 in the full dataset. Box-whisker plots similarly show the distribution across 100 bootstrap replicates. (C) Receiver operator characteristic (ROC) curves indicate the sensitivity and specificity of DE gene classification at varying significance thresholds. For each sample size, the filled line represents the median ROC sensitivity amongst 100 bootstrap replicates, while the dashed lines represent the 0.05 and 0.95 quantiles. (D) The ability to significantly detect small effect size DE genes increases as sample size increases. At each indicated subsample size, box-whisker plot (displaying 0.05, 0.25, 0.5, 0.75, 0.95 quantiles) indicates effect size of true significant DE genes (The effect size and true classifications of significant DE genes are defined as those at FDR<0.01 with the full dataset). (E) Winner’s curse effects decrease with increasing sample size. The median difference between the estimated effect size of DE genes and the true effect sizes estimated from the full dataset. Box-whisker plots depict the 0.05, 0.25, 0.5, 0.75, and 0.95 quantiles among 100 bootstrap replicates. (F-L) Same as (A-E) but each sample was subsampled at the level of RNA-seq reads to a depth of 25 million mapped reads. (K-O) Same as (A-E) but each sample was subsampled to 10 million mapped reads.

### Characterizing variability in gene expression

The main focus of our study was to assess and compare the inter-individual variability in gene expression between the two species. To do so, we estimated overdispersion for each gene in chimpanzees and humans separately (see Methods). Briefly, we assumed that the measurement process is captured by Poisson sampling, and that true gene expression follows a Gamma distribution. These assumptions are supported by theory (Pachter, 2011) and empirical data (Marioni et al., 2008). Accordingly, we used a negative binomial regression model to fit the RNA-seq data (22,23). Fitting the model to the data from each gene yields estimates of the mean and biological variance of gene expression, which result in the overdispersion of the observed read counts relative to a Poisson distribution.

Consistent with previous findings (BLUEPRINT Consortium et al., 2017; Eling et al., 2018; Robinson et al., 2010), we observed that overdispersion is strongly correlated with mean expression, due to reasons which may be technical or biological in nature. To understand the properties of gene expression variability that are independent of mean gene expression level, we regressed out the mean from the overdispersion estimates, to obtain a mean-corrected value, hereafter referred to as “dispersion” (Fig1A,B). A dispersion greater than 0 indicates more variability than expected given the gene’s expression level. Conceptually, our approach is similar to a method recently devised to identify differentially dispersed genes using single cell gene expression data (Eling et al., 2018). Using this approach, one can identify genes whose inter-individual variance in expression is different, irrespective of their mean or median expression levels. For example, while the genes *TERF2* and *SNORD14E* have similar median expression levels, *TERF2* has a lower dispersion than *SNORD14E*, in both species (Fig1B). We used a bootstrap test (Methods) to assess the stability of dispersion estimates and identified genes whose estimated dispersion is significantly different between species. For example, *ZNF514* has higher dispersion in humans than chimpanzees, despite being similarly expressed in both species (Fig1B). In total, we identified 2,658 inter-species differentially dispersed genes (FDR<0.1). Inter-species differences in mean expression levels cannot explain this finding, as differentially dispersed genes are generally not differentially expressed (FigS4).

**Fig1.**
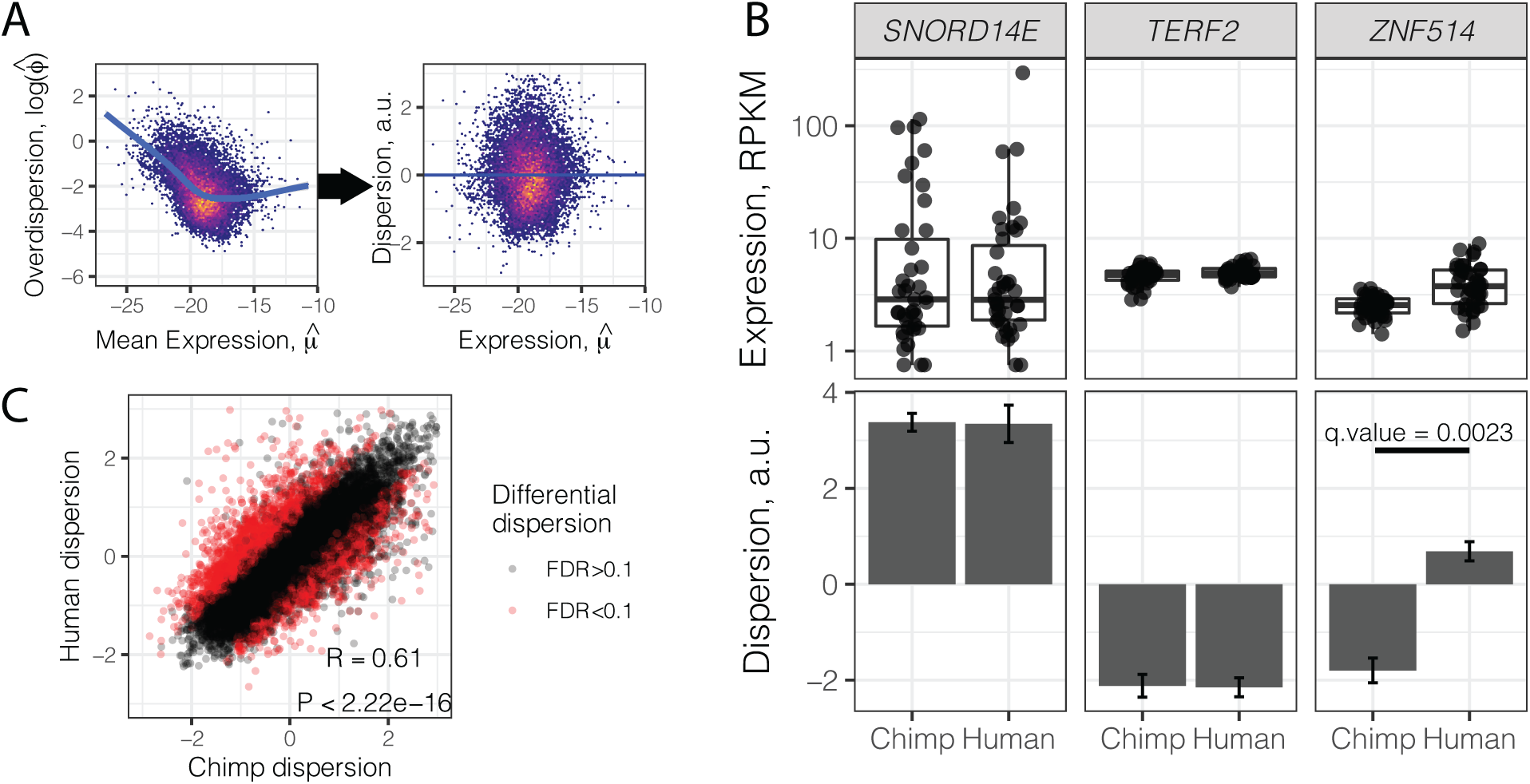
Gene variance independent of expression is correlated across species. (A) To estimate dispersion, RNA-seq counts for 39 human heart tissue samples were used to estimate gene-wise mean (μ) and overdispersion (ϕ) parameters. Across all genes, overdispersion is correlated with mean expression (left) in the hexbin scatter plot. We regressed out this correlation, using the residual of a LOESS fitted line (blue) as a metric (dispersion) of the variability of a gene’s expression across a population relative to similarly expressed genes. (B) Dispersion estimates and the underlying expression in each sample for 3 similarly expressed genes in human and chimpanzee. Error bars represent bootstrapped standard error. Q-value for *ZNF514* represents an estimate of FDR after genome-wide multiple hypothesis testing correction. (C) Dispersion estimates across all genes are correlated across human and chimpanzee, despite identification of thousands of differentially dispersed genes in red. R and P-value correspond to Pearson’s correlation.

**FigS4.**
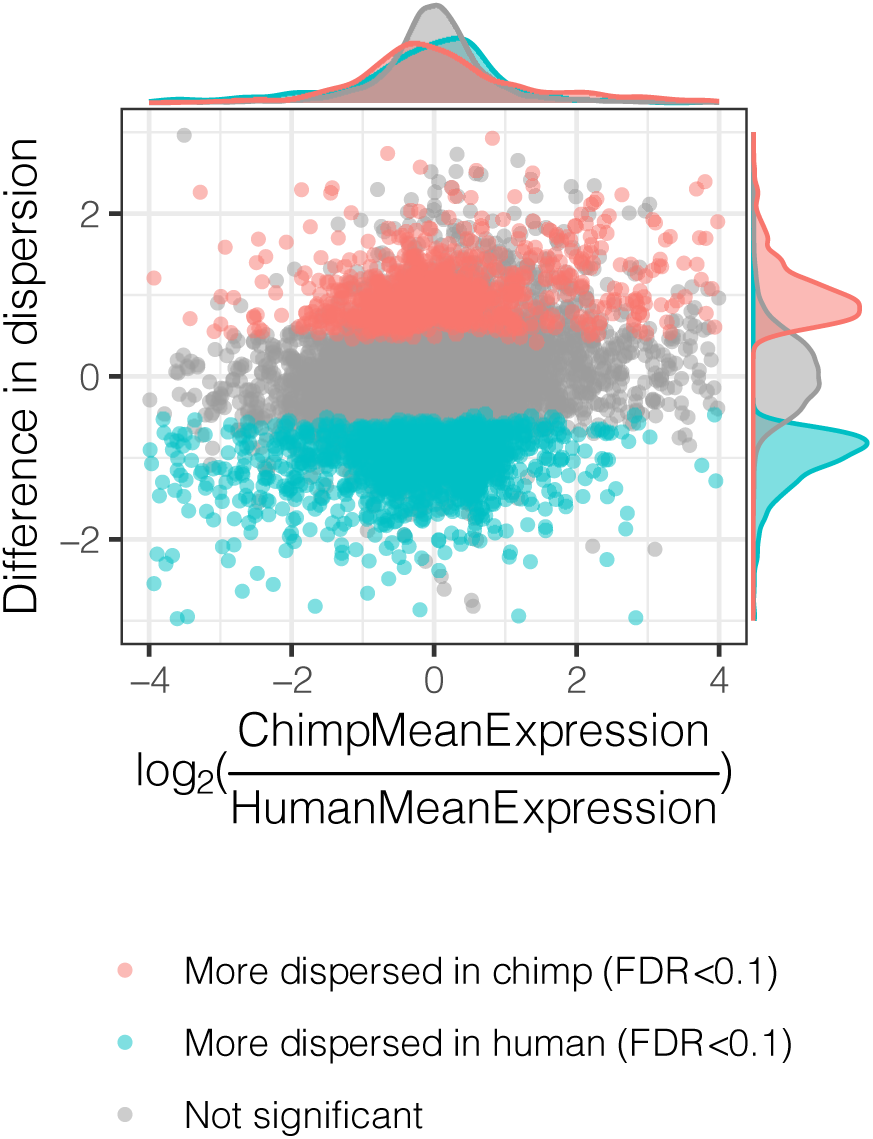
Interspecies dispersion estimates are largely independent from interspecies mean expression changes. Across all genes, the interspecies difference in dispersion, colored by significance, is plotted versus difference in mean expression.

Despite the identification of thousands of differentially dispersed genes between the species, gene-wise dispersion estimates are well correlated in humans and chimpanzees overall (R=0.60, Fig1C, TableS3), suggesting similar determinants of variability in the two species. We asked what other gene properties may be associated with the degree of gene expression variability. Specifically, we hypothesized that highly conserved essential genes, whose coding regions evolve under strong negative selection, would also be less variable in their expression across individuals. Indeed, when we ranked and grouped genes by degree of protein coding conservation (assessed by percent amino acid identity between human and chimpanzee; Fig2A), or by the ratio of nonsynonymous to synonymous codon changes (dN/dS) across mammals (Fig2B), we found that lower dispersion in expression levels is associated with higher protein coding conservation. Using the prevalence of genetic diversity data from humans, we then categorized genes as loss-of-function-tolerant (ExAC pLI score < 0.1) or loss-of-function-intolerant (ExAC pLI score > 0.9) (Exome Aggregation Consortium et al., 2016). We found that the expression of loss-of-function-intolerant genes is associated with lower dispersion (Fig2C). Unsurprisingly, these features of low dispersion genes similarly hold in chimpanzee (FigS5) as in human (Fig2).

**Fig2.**
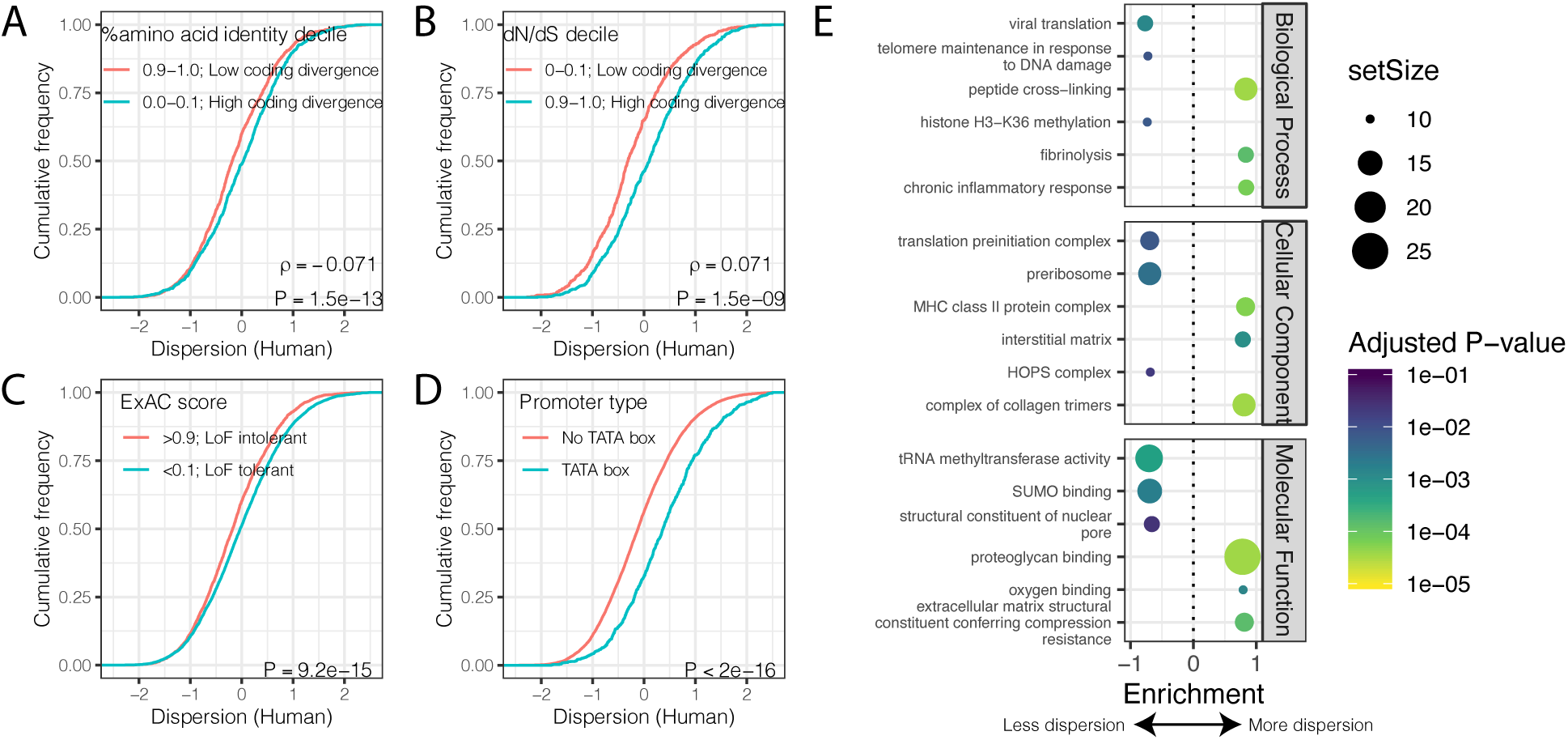
Gene features correlated with expression variability. (A) Protein coding genes with high coding divergence (defined by amino acid identity between chimpanzee and human) generally have higher variability than genes with low coding divergence. The distribution of dispersion estimates is plotted as the empirical cumulative distribution function (ECDF) for the top and bottom decile genes by percent identity. (B) Same as (A) but defining coding divergence based on rate of non-synonymous to synonymous substitution rates (dN/dS) across mammals. (C) Loss-of-function tolerant (LoF tolerant) genes, defined by pLI score (Exome Aggregation Consortium et al., 2016), generally have higher variability than loss-of-function intolerant (LoF intolerant) genes. (D) TATA box genes generally show higher variability. P-values and ρ correlation coefficient provided for (A) and (B) represent Spearman correlation across all quantiles, rather than just the upper and lower decile, which are plotted for similar visual interpretation as (C) and (D), where the P-values provided represent a two-sided Mann-Whitney U-test. (E) Gene set enrichment analysis of genes ordered by human dispersion estimates. Only the top and bottom 3 most enriched significant categories (Adjusted P-value < 0.05) are shown for each ontology set for space.

**FigS5.**
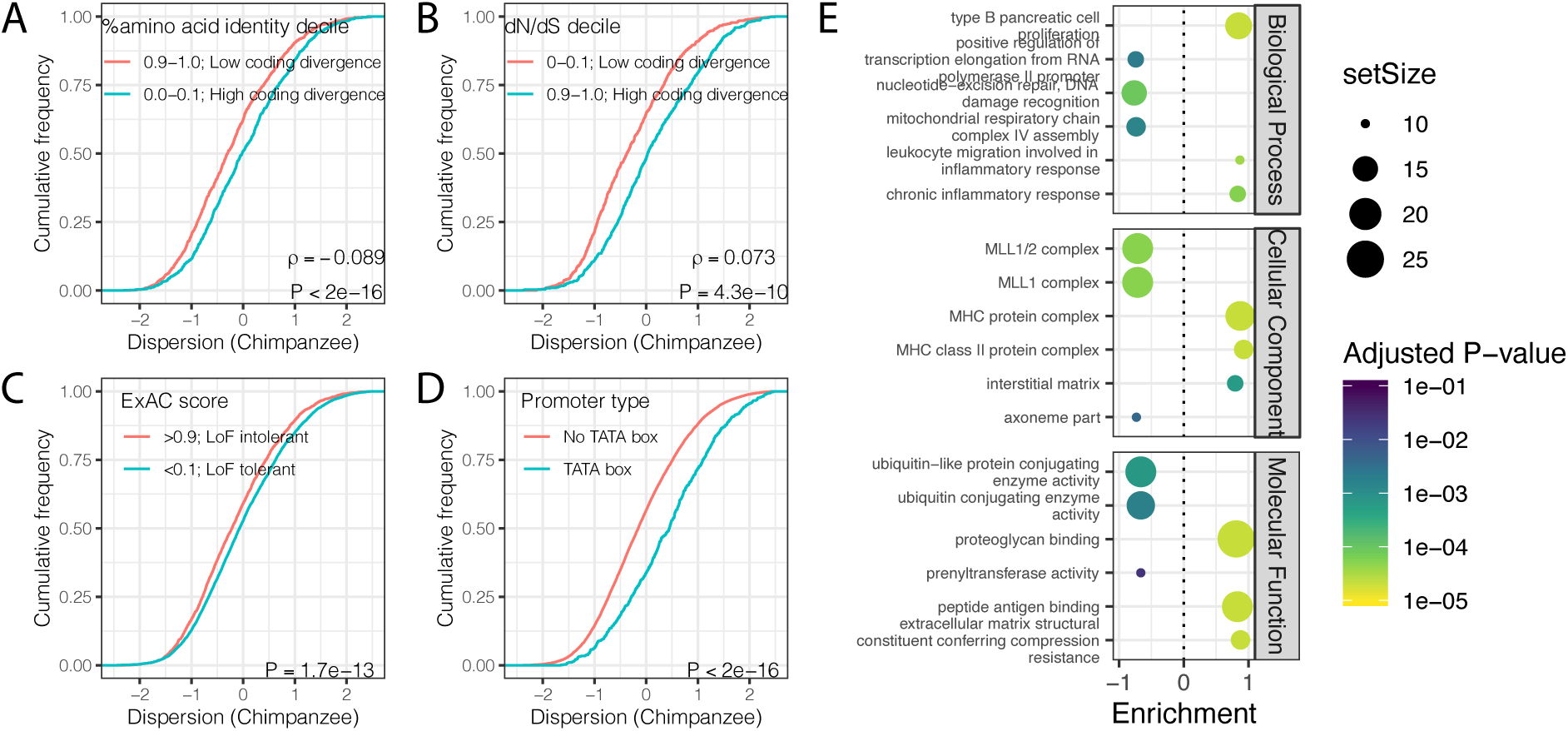
Gene features correlated with expression variability (chimpanzee) (A) Protein coding genes with high coding divergence (defined by amino acid identity between chimpanzee and human) generally have higher variability than genes with low coding divergence. The distribution of chimpanzee dispersion estimates is plotted as the empirical cumulative distribution function (ECDF) for the top and bottom decile genes by percent identity. (B) Same as (A) but defining coding divergence based on rate of non-synonymous to synonymous substitution rates (dN/dS) across mammals. (C) Loss-of-function tolerant (LoF tolerant) genes, defined by pLI score (Exome Aggregation Consortium et al., 2016), generally have higher variability than loss-of-function intolerant (LoF intolerant) genes. (D) TATA box genes generally show higher variability. P-values and ρ correlation coefficient provided for (A) and (B) represent Spearman correlation across all quantiles, rather than just the upper and lower decile, which are plotted for similar visual interpretation as (C) and (D), where the P-values provided represent a two-sided Mann-Whitney U-test. (E) Gene set enrichment analysis of genes ordered by chimpanzee dispersion estimates. Only the top and bottom 3 most enriched significant categories (Adjusted P-value < 0.05) are shown for each ontology set for space.

Next, we asked what functional Gene Ontology (GO) categories are enriched among the most or least dispersed genes. To do this, we ordered genes by their dispersion estimate in humans (Fig2E) or chimpanzees (FigS5E) and used gene set enrichment analysis (GSEA) (Subramanian et al., 2005) to identify GO categories enriched at the top or bottom of the list. We found that genes with housekeeping functions universal to all cell-types, such as genes related to transcription initiation and tRNA modification, are among the most enriched gene categories associated with low dispersion (Fig2E, FigS5E, TableS4). Conversely, genes with high dispersion are enriched for categories like fibrinolysis (the breakdown of blood clots), oxygen sensing, and other cardiovascular related functions (Fig2E, FigS5E, TableS4). Finally, consistent with previous reports (de Jong et al., 2019; Hagai et al., 2018; Ravarani et al., 2016), we found that genes with TATA boxes are associated with higher dispersion (Fig2D), potentially because TATA boxes are associated with higher transcriptional noise at the molecular level (Blake et al., 2006; Raser, 2004; Ravarani et al., 2016). However, a possible technical explanation for this observation is that TATA box genes are enriched among cell-type-specific genes (Schug et al., 2005) and cell-type heterogeneity between individuals or samples could contribute to the observed dispersion.

To investigate the possibility that differences in cell composition between samples contribute to our observations of gene expression dispersion, we first asked whether dispersion is associated with the degree of cell-type-specificity of expression. We hypothesized that genes with high inter-individual dispersion are more likely to have cell-type-specific gene expression signatures in single cell RNA-seq datasets of heart tissue. To examine this we turned to the *Tabula Muris* dataset, a comprehensive single cell transcriptomics dataset which includes single cell RNA-seq data from adult mouse heart tissue (The Tabula Muris Consortium et al., 2018). Qualitatively, we observed that the most highly dispersed genes in our bulk chimpanzee and human samples have more cell-type-specific expression among the 9 heart cell types identified in *Tabula Muris* (Fig3A). Conversely, the lowest dispersed genes are more evenly expressed across cell types (Fig3B). More generally, when we summarized the level of cell-type-specific expression for each gene as a single summary statistic, τ (Kryuchkova-Mostacci and Robinson-Rechavi, 2016; Yanai et al., 2005), we found that dispersion is strongly correlated with cell-type-specificity (R=0.35, *P* < 2×10^−16^; Fig3C).

**Fig3.**
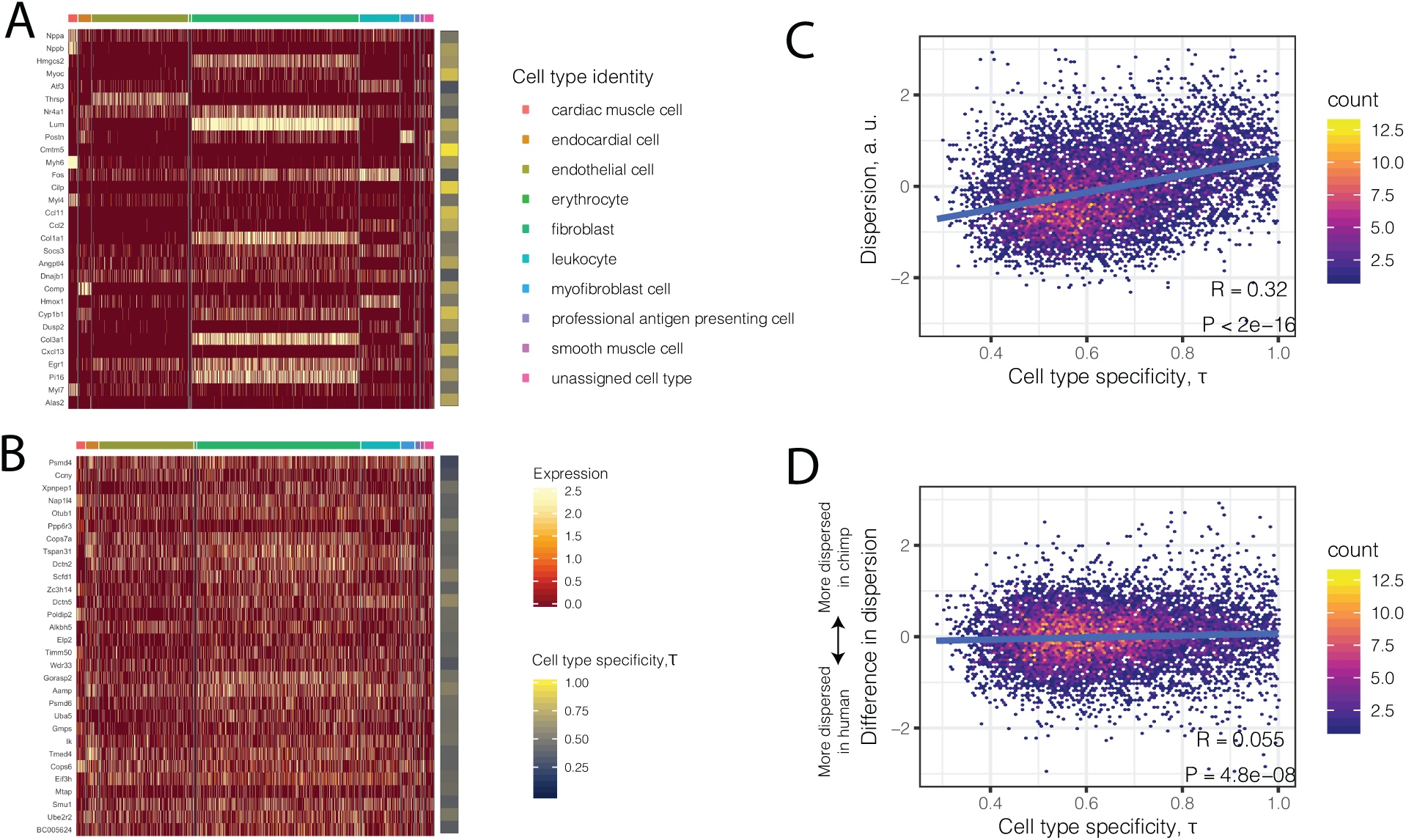
High dispersion genes are expressed in a cell-type-specific manner. (A) The top 30 most dispersed genes (the mean of the dispersion estimates between human and chimpanzee) are shown in a mouse heart single cell RNA-seq dataset (The Tabula Muris Consortium et al., 2018). Each row is a gene. Each column is a single cell, grouped by cell type (colors at top of columns). Normalized expression is colored in the body of the heatmap. The τ statistic, colored on the right of the heatmap, summarizes for each gene how cell-type-specific the gene is, ranging from 0 (equally expressed among all cell types) to 1 (expressed exclusively in a single cell type). (B) The same as (A) but for the bottom 30 most dispersed genes. (C) Across all genes, a hexbin scatterplot shows the correlation between cell-type-specificity (τ) estimated from mouse single cell RNA-seq data, and dispersion (mean of human and chimpanzee dispersion for each gene) estimated from the bulk RNA-seq data. (D) The difference in dispersion between chimpanzee and human is only weakly correlated with cell-type-specificity. R and P-value for (C) and (D) represent Pearson’s correlation.

Given the correlation between dispersion and the extent of cell-type-specific regulation, we sought to estimate the proportions of different cell types amongst our bulk RNA-seq samples for both chimpanzee and human. We applied CIBERSORT (Newman et al., 2015) to the bulk RNA-seq profiles using reference cell type profiles derived from *Tabula Muris* (Methods). As expected, cell type proportions between chimpanzee and human hearts are qualitatively similar, though much inter-individual variation exists in both species for particular cell types, such as cardiac muscle cells and myofibroblasts (FigS6). Put together, these analyses indicate that the levels of inter-individual dispersion we observed in humans and chimpanzees, and the high similarity in dispersion observed across species, are strongly driven by cell type heterogeneity between samples. These cell type composition differences may be partly due to both technical differences between sample preparations, as well as biological differences between individuals. Importantly, however, differences in dispersion between chimpanzee and human is much less likely to be driven by cell type heterogeneity, though there is a relatively small but significant correlation with τ (R=0.07, *P* = 2.2×10^−12^; Fig3D).

**FigS6.**
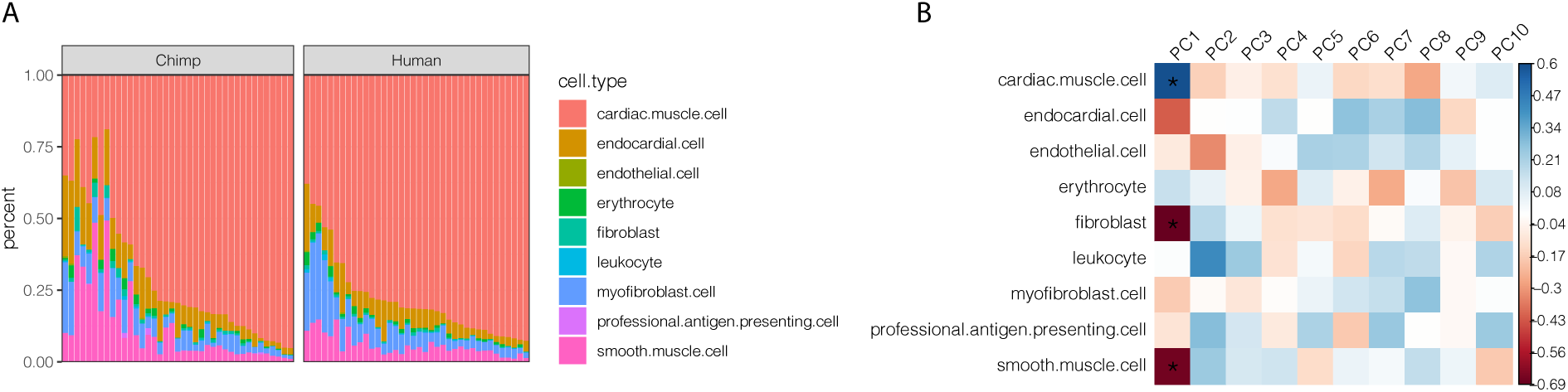
Cell composition varies across individuals but is captured by PCs included in eQTL mapping. (A) The cell type composition estimates of chimpanzee and human bulk RNA-seq samples used in dispersion. Height of colored bars represents estimates of the proportion of each heart cell type. (B) A heatmap of the Pearson correlation between the principle components of the chimpanzee samples used in eQTL mapping, to their cell composition estimates. Principle component 1 corelates with cell type composition (*Benjamini-Hochberg corrected P<0.005).

Having more confidence in the observation of inter-species differences in dispersion, we asked what gene categories are enriched among differentially dispersed genes. We performed GSEA (Subramanian et al., 2005), ranking genes by the polarized significance level of the chimpanzee-human difference in dispersion estimates. We found that genes more variable in human are enriched for mitotic regulators (FigS7A). Given evidence that ischemia may induce mitosis in adult mammalian cardiac cells (Kajstura et al., 1998; Kimura et al., 2017; Nakada et al., 2017), the enrichment for mitotic regulators may reflect the highly variable life histories in GTEx samples, a large fraction of which are sourced from organ donors with ischemic cardiovascular disease. Conversely, we found that genes related to immune function are more variable in chimpanzee. We note that some of our chimpanzee individuals were sourced from laboratory settings and have been challenged with HBV or HCV viral infections. However, our GSEA results are robust to the exclusion of these samples (TableS5, TableS6).

**FigS7.**
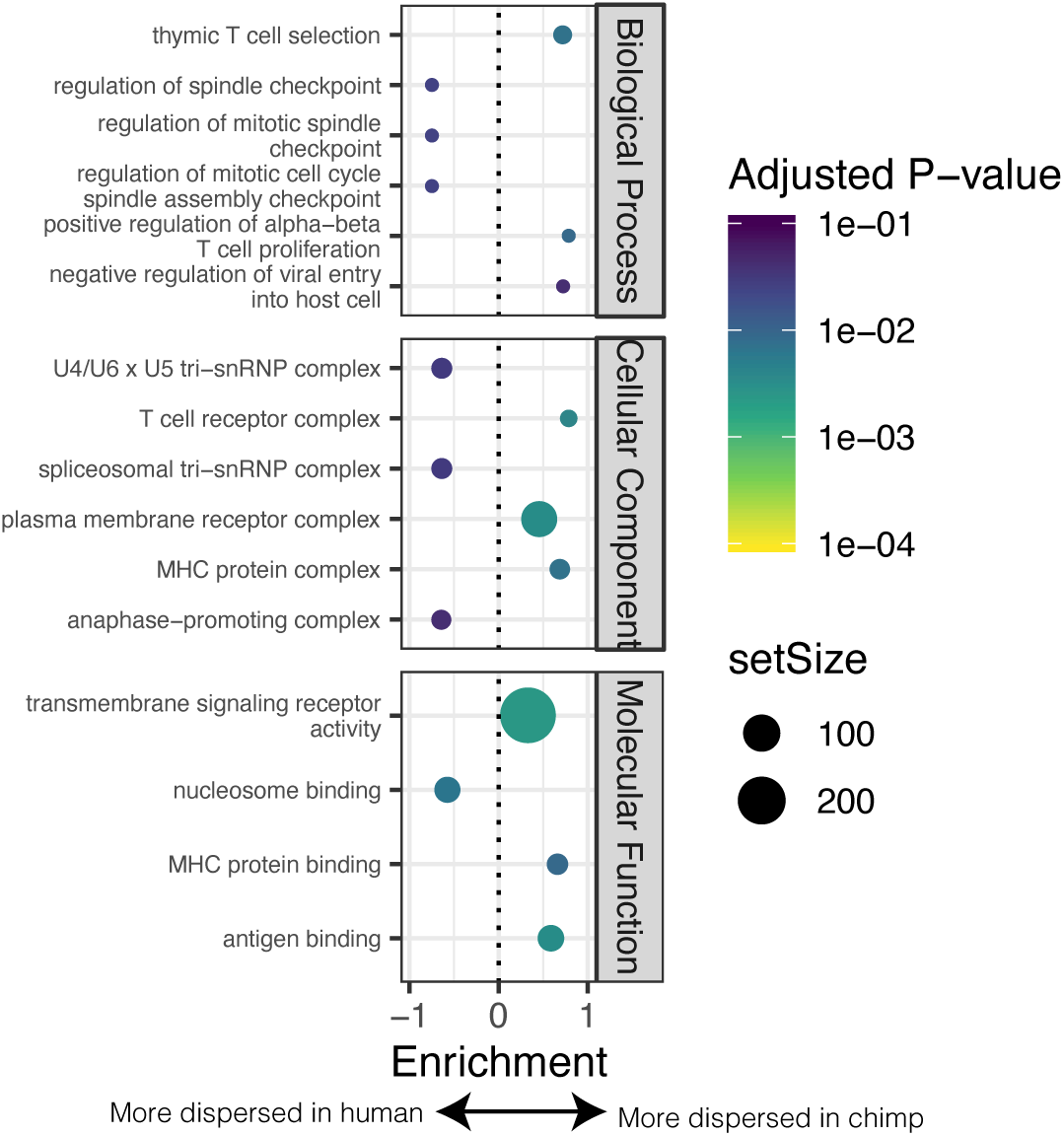
GSEA of genes with different levels of dispersion between species. (A) GSEA of genes ordered by difference in dispersion estimates. Genes with more variable expression in chimpanzee often relate to immune response ontology terms. Genes with more variable expression in human often relate to heart function related ontology terms. Only the top 3 and bottom 3 most enriched significant terms for each sub ontology category are shown for space.

### Within-species genetic variation contributes to inter-species differences in variability

Although potentially technical in nature, inter-individual differences in cellular composition provide a partial explanation for our observation of similar dispersion estimates across species. We looked for evidence that genetic diversity also drives dispersion. We asked whether inter-species differences in dispersion, which are much less likely to be explained by cellular composition, are associated with corresponding differences in selection pressures between the species. We reasoned that if inter-species differences in dispersion are partially driven by inter-species differences in selection pressures, we may see differences between the species in genetic signatures that are consistent with natural selection near the differentially dispersed genes. More specifically, we expect that genes with particularly low dispersion in human compared to chimpanzee will also display more constraint at the coding level in human than in chimpanzee. To assess this, we analyzed genotype data from our chimpanzee and human cohorts. We obtained human genotype data from the GTEx consortium, and chimpanzee data by performing high-coverage whole genome sequencing on the 39 chimpanzee samples used in this study (>30X genome coverage obtained in all samples, see tableS7); these data represent roughly a 50% increase in the number of high-coverage *Pan troglodytes* genomes currently available (de Manuel et al., 2016). We used the sequencing data to identify nearly 2.9 million novel chimpanzee SNPs with a minor allele frequency (MAF) greater than 10%.

Using the genotype data, we asked if differences in expression dispersion are associated with differences in evolutionary constraint on protein coding regions in the human and chimpanzee lineages. For each gene, we calculated the ratio of non-synonymous polymorphisms (P_n_) scaled to synonymous polymorphisms (P_s_) within each species. This P_n_/P_s_ metric may be used to assess purifying and diversifying selection pressures acting on coding sequences *within* species (Fuller et al., 2015; Huguet et al., 2014; Tanaka and Nei, 1989), which thus may correspond to our measurements of within-species expression dispersion. In contrast, the more often-utilized dN/dS metric is based on fixed coding differences between species and therefore not well suited to identify the signatures of selection that uniquely confine variability *within* a species. As expected, we found that loss-of-function tolerant genes have higher P_n_/P_s_ than loss- of-function intolerant genes (Fig4A). When we compared P_n_/P_s_ between humans and chimpanzees, we found that the inter-species ratio of P_n_/P_s_ is positively correlated with the difference in gene expression dispersion between species (Fig4B). That is, on average, genes with higher dispersion in chimpanzees than in humans have a higher abundance of non-synonymous polymorphisms (P_n_/P_s_) in chimpanzees compared to the orthologous genes in humans. This suggests that inter-species differences in selection pressures at polymorphic loci play a role in the observed inter-species differences in expression dispersion.

**Fig4.**
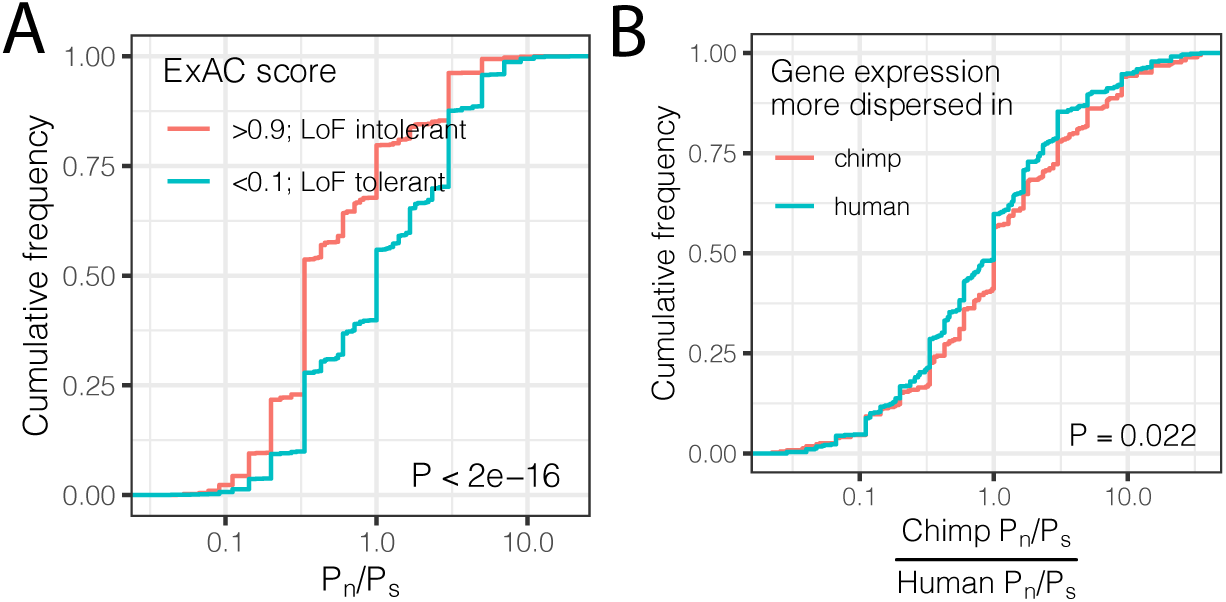
Interspecies differences in dispersion correlate to interspecies differences in coding constraint. (A) The number of nonsynonymous polymorphisms scaled to synonymous polymorphisms (P_n_/P_s_) for each gene was calculated in GTEx human population. Loss-of-function tolerant (LoF intolerant) genes, defined by pLI score (Exome Aggregation Consortium et al., 2016), generally have higher P_n_/P_s_ than loss of function tolerant (LoF tolerant) genes, as shown in ECDF plot. (B) P_n_/P_s_ was calculated for both human and chimpanzee. The distribution of the chimpanzee P_n_/P_s_ to human P_n_/P_s_ ratio is plotted as an ECDF, grouped by whether the gene has higher dispersion estimate in chimpanzee than in human. P-value indicates a Mann Whitney U-test.

### Genes with eQTLs are shared across species more often than expected by chance

The correlation between the degree of evolutionary constraint on coding sequences and dispersion of expression at the gene level suggests that differences in cellular composition are not the only explanation for differences in dispersion. Motivated by this notion, we searched for further evidence for genetic regulation of dispersion by identifying genes associated with eQTLs - eGenes - in both humans and chimpanzees. In humans, we obtained a list of 11,682 heart left ventricle eGenes identified by the GTEx consortium (with a sample size of 386 individuals). In chimpanzee, we used the genotype and expression data we collected to map *cis* eQTL SNPs within 250kb of each of the 13,545 expressed genes. We included 10 principal components as covariates (FigS8A) and accounted for genetic relatedness between individuals and population structure, which we inferred from the genotype data (Methods and FigS9). Because gene expression principle components are correlated with cellular heterogeneity (FigS5B), their inclusion as covariates in the eQTL linear modeling helps correct for this heterogeneity and additional unobserved technical sources of variation. Using this approach, we identified 310 eGenes in chimpanzee hearts (FDR<0.1; FigS8A-B, tableS8). Consistent with previous eQTL studies in primates (Jasinska et al., 2017; Pickrell et al., 2010; Tung et al., 2015; Veyrieras et al., 2008), we found that the chimpanzee eQTL SNPs are enriched near transcription start sites (FigS8C).

**FigS8.**
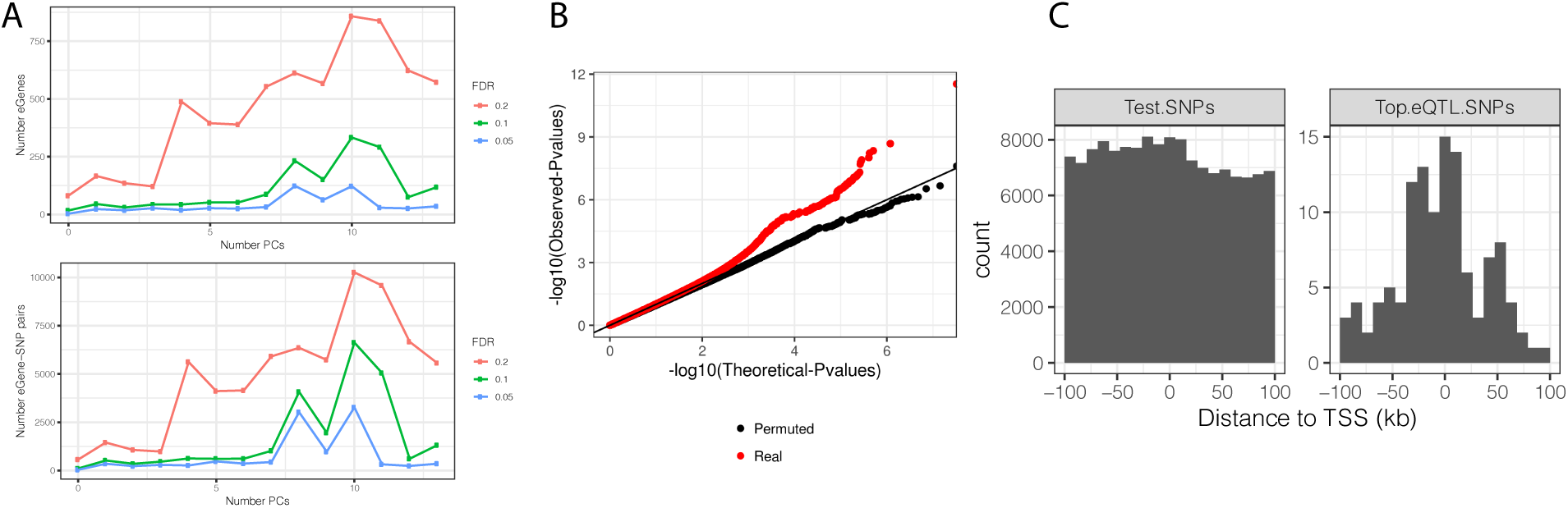
(A) The number of significant variant:gene pairs and eGenes is plotted when varying numbers of principle components are included as covariates. (B) QQ-plot of P-values for all variant:gene pairs tested shows inflation compared to sample permutation control. (C) For eGenes (FDR<0.1), the distribution of distances of the top eQTL variant to the transcription start site, compared to the distribution of all test variants in eGenes.

**FigS9.**
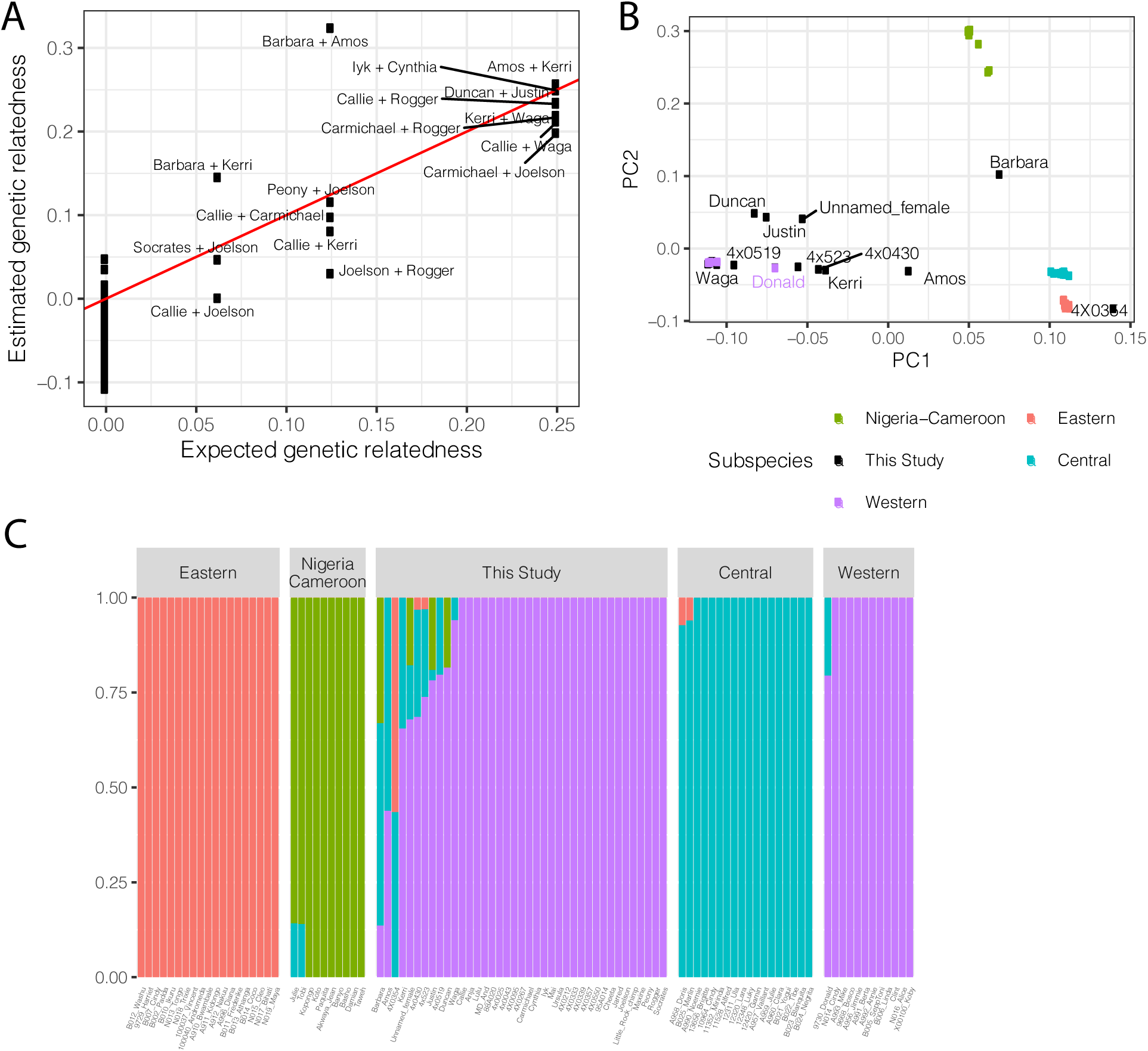
Population structure and relatedness of chimpanzee cohort based on whole genome sequencing genotyping. (A) Scatter plot of the expected and observed genetic relatedness of each pair of chimpanzees for which we have pedigree information. We expected 8 first degree relationships (relatedness=0.25), 5 second degree relationships, and 3 third degree relationships. (B) The chimpanzees sequenced in this study are generally Western chimpanzees, though recent admixture with other subspecies is prevalent for many of these captive-born chimpanzees. Principle component analysis of genotypes of newly sequenced and previously sequenced (de Manuel et al., 2016) chimpanzees. Individuals colored by subspecies for previously sequenced chimpanzees (Eastern, *Pan troglodytes schweinfurthii*; Western, *Pan troglodytes verus;* Central, *Pan troglodytes troglodytes;* Nigeria-Cameroon, *Pan troglodytes ellioti*). Donald is a previously sequenced captive born chimpanzee with known Western-Central admixture (de Manuel et al., 2016). Only newly sequenced chimpanzees with signs of admixture are labelled. (C) Admixture analysis (K=4) of newly sequenced and previously sequenced chimpanzees.

We considered the overlap of eGenes in humans and chimpanzees. When considering only one-to-one orthologs tested for eGenes in both species, there is no significant overlap of eGenes in the two species (Odds Ratio=1.03, *P*=0.46, hypergeometric test, FigS10A). However, this comparison is affected by the substantial difference in power to detect eQTLs between the large human sample used in GTEx and the relatively small chimpanzee sample we collected. To address this, we iteratively subsampled the human cohort to sample sizes comparable to our chimpanzee cohort, and re-mapped human eQTLs. When we compared lists of eGenes identified in humans and chimpanzees using similar sample sizes, we found a greater overlap of eGenes in the two species than expected by chance (FigS10B). This observation is robust even if we use the eQTL results of the full GTEx dataset, as long as we perform the comparison by using the largest-effect eQTLs (those that can be identified as significant even in smaller sample sizes). Specifically, of the top 500 significant GTEx eGenes (Fig5A, FigS10C), 21 are also found to be eGenes in chimpanzee (FDR < 0.1), a significant enrichment (Odds Ratio=2.05, *P*=0.003, hypergeometric test). Importantly, we found that species-specific eGenes have higher species-specific dispersion (Fig5B). That is, eGenes identified in chimpanzees but not humans have higher dispersion in chimpanzees and vice versa (*P*=3.7×10^−11^; Fig5B). This observation is consistent with a genetic contribution to inter-species differences in gene expression dispersion. Furthermore, the observation that more eGenes are shared among humans and chimpanzees than expected by chance suggests that the regulation of these genes evolves under less evolutionary constraint.

**FigS10.**
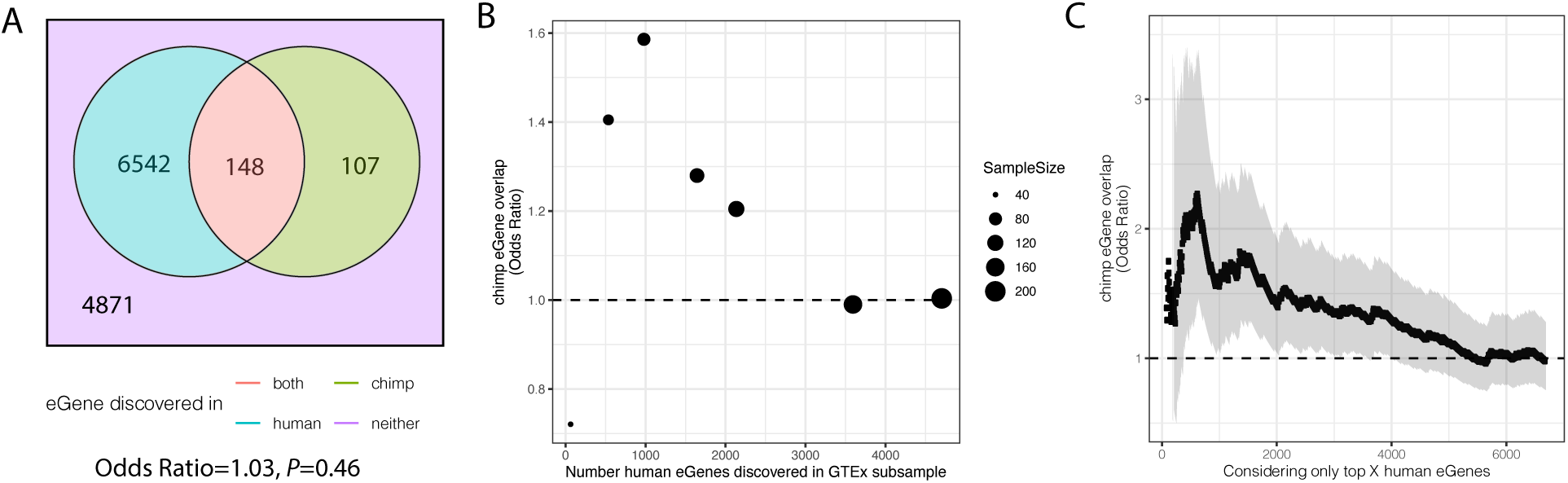
Significant overlap of eGenes between chimpanzee and human is observed when studies are similarly powered. (A). Overlap of eGenes identified in chimpanzee (FDR<0.1, n=38) and human (FDR<0.1, sourced from GTEx; n=386). Only genes that are one-to-one orthologs and passed expression filters for testing were considered. (B) The overlap of eGenes is more than expected by chance (assessed by odds ratio > 1) when the overpowered GTEx dataset is randomly subsampled to sizes more comparable to the chimpanzee dataset and genes are retested for eGene activity in the subsample. (C) Similar to (B) when only the top X genes (ranked by FDR using the full GTEx dataset) are considered human eGenes, the overlap becomes significant as eGene classification becomes more stringent. Shaded region represents 95% confidence interval of odds ratio.

**Fig5.**
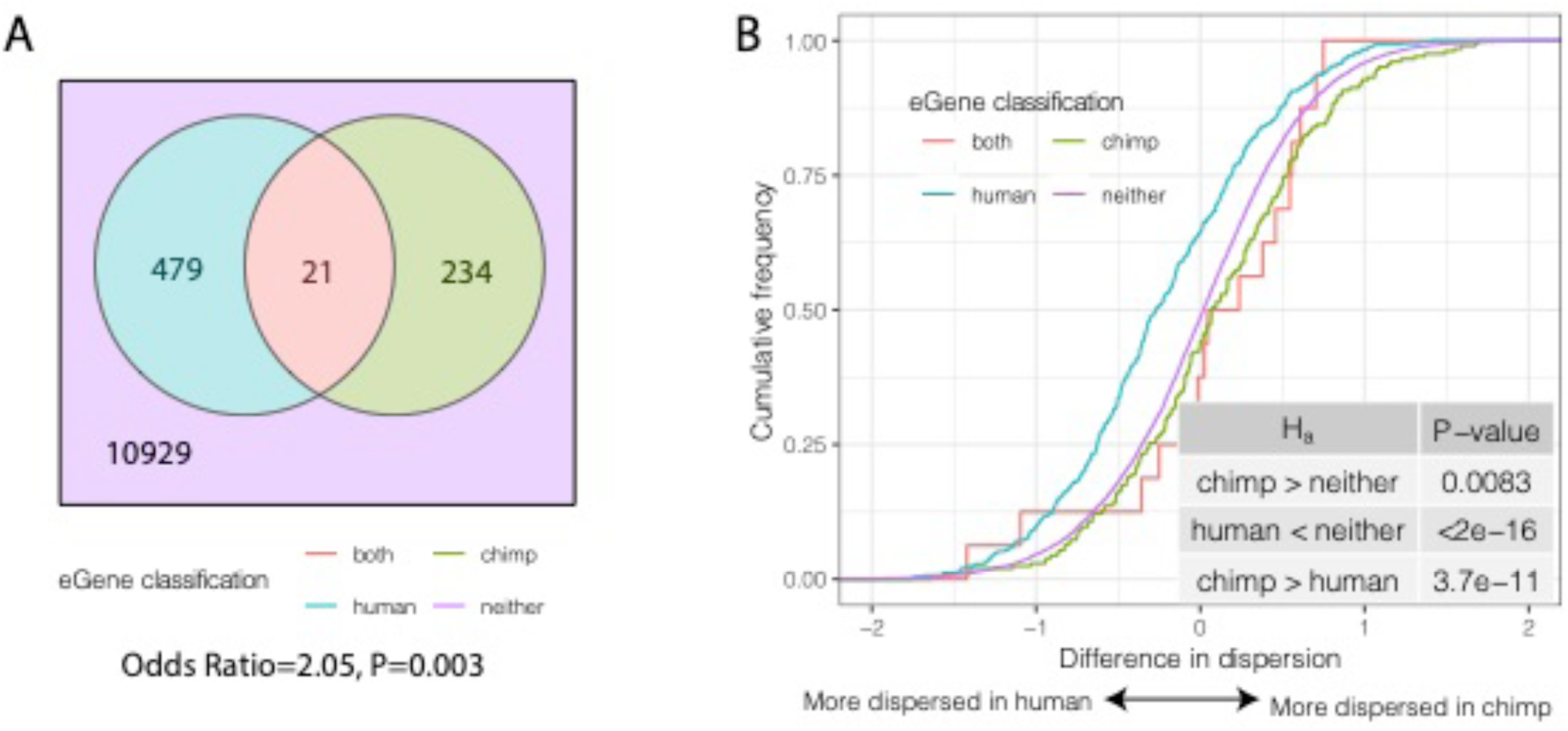
Species-sharing and dispersion of eGenes. (A). eGenes were classified by a 10% FDR threshold in chimpanzee and considering only the top 500 eGenes by FDR in human (GTEx). (B) ECDF of the difference in dispersion of genes between chimpanzee and human. Chimpanzee-specific eGenes are more dispersed in chimpanzee; human-specific eGenes are more dispersed in human. P-values provided for one-sided Mann-Whitney U-test with the noted alternative hypothesis.

### Shared and species-specific eGenes may evolve under different selection pressures

To further examine whether shared and species-specific eGenes may evolve under different selection pressures than non-eGenes, we examined other indicators of selection for each of these eGene groups. We found that eGenes have higher inter-species differences in expression levels than non-eGenes (Fig6A), and eGenes identified in both species have even larger differences in mean expression levels between species than eGenes identified in only in chimpanzees or humans. These observations are consistent with the notion that the regulation of genes associated with eQTLs tend to evolve under less evolutionary constraint. Furthermore, eGenes tend to have lower levels of coding conservation in both species, as measured by amino acid identity between human and chimpanzee (Fig6B) or dn/ds across mammals (Fig6C).

**Fig6.**
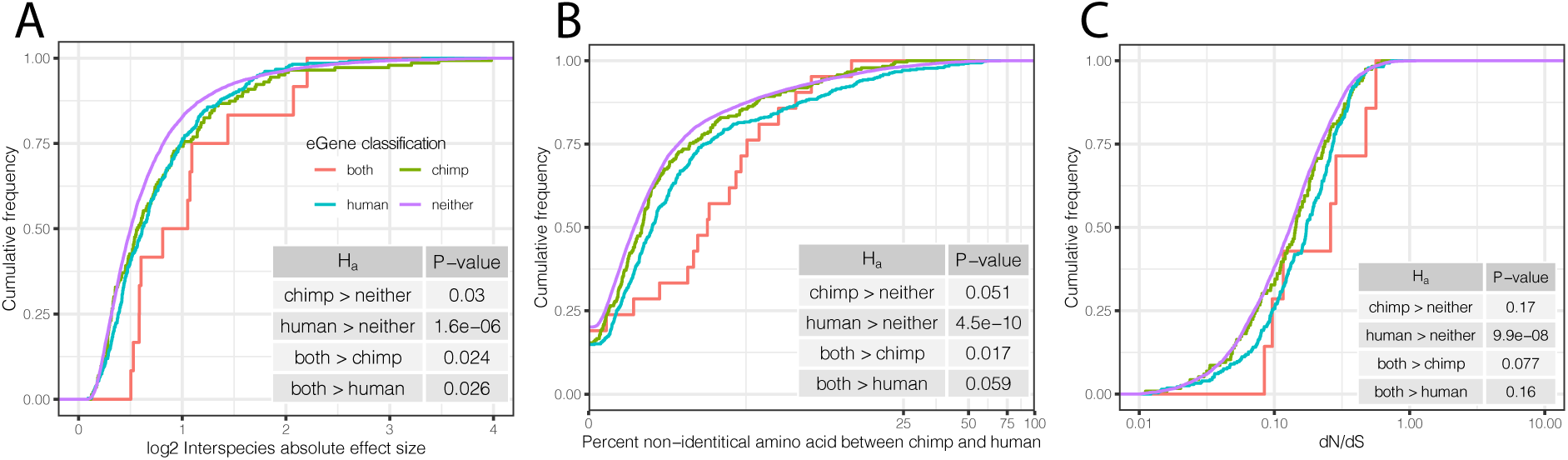
Characteristics of eGenes are consistent with less constraint on eGene expression. (A) eGenes are more differentially expressed between species than non-eGenes, with eGenes detected in both species being even more differentially expressed. The distribution of the inter-species differential expression effect size is plotted for each eGene group as an ECDF. (B) eGenes are more diverged at amino acid level than non-eGenes. (C) Human-specific eGenes and shared eGenes are more divergent than expected under neutrality. The analogous test for chimpanzee-specific eGenes displayed a shift that was not statistically significant, though our classification of eGenes in chimpanzee may be underpowered. P-values provided for one-sided Mann-Whitney U-tests with the noted alternative hypothesis.

We next performed GO enrichment analysis (hypergeometric test) to ask which functional classes are identified as eGenes in both human and chimpanzee. We reasoned that the 21 genes identified as shared eGenes may have high levels of genetically regulated variability, which may indicate expression evolution at a neutral rate or faster. We found that these genes are strongly enriched for immune response genes, including major histocompatibility complex (MHC) genes (FigS10A, TableS9). This observation is consistent with previous reports that immune genes evolve under strong directional and balancing selection pressures across vertebrates, in part to respond to ever-evolving pathogen challenges (Ejsmond and Radwan, 2015; Hagai et al., 2018; Lam et al., 2017; Shultz and Sackton, 2019).

Given the evidence that highly dispersed genes and eGenes are associated with relaxed evolutionary constraint, we next asked which gene classes are enriched among chimpanzee-specific eGenes. This set of genes may be subject to stronger stabilizing selection in the human lineage. We approached this question by considering all eGenes discovered in the full GTEx dataset (FDR<0.1) as human eGenes, as this is the most stringent way to classify eGenes as chimpanzee-specific. We identified 148 chimpanzee-specific eGenes, which we found to be significantly enriched for transcriptional regulation terms (FigS10B, TableS10).

**FigS10.**
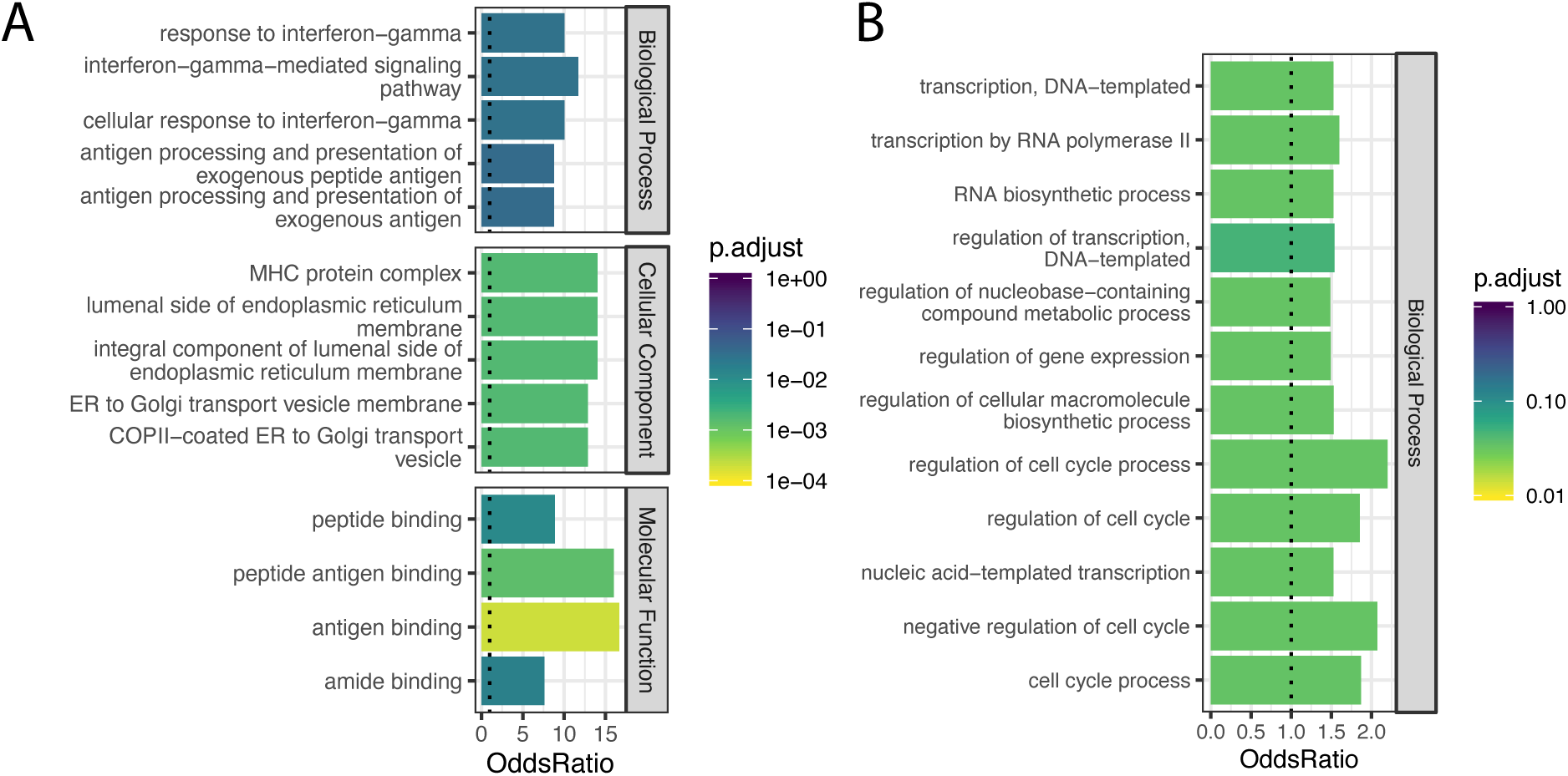
Gene categories of human-chimpanzee shared and chimpanzee-specific eGenes. (A) Gene ontology categories enriched amongst the 21 eGenes identified in both species (foreground), compared to the 734 eGenes identified in only chimpanzee or only human (universe/background). Among the significant categories (Adjusted P-value<0.05), only the top 5 most enriched gene categories for each ontology set are shown for space. (B) Gene ontology categories enriched amongst the 148 eGenes identified only in chimpanzee, compared all 6797 eGenes identified in either species using the full GTEx dataset (universe/background). All significant categories are shown.

### Effects of trans-species polymorphisms on gene expression

Genetically driven variability in gene expression may also arise due to overdominant or frequency-dependent selection on gene regulation, which maintains polymorphisms over evolutionary time through balancing selection (Croze et al., 2016; Těšický and Vinkler, 2015). MHC and other immune genes are well known targets of these modes of selection, as host immune systems are under constant evolutionary pressure to diversify in response to quickly evolving pathogens. As such, these genes sometimes contain trans-species polymorphisms maintained through evolutionary time by balancing selection (Croze et al., 2016; Těšický and Vinkler, 2015).

**FigS11.**
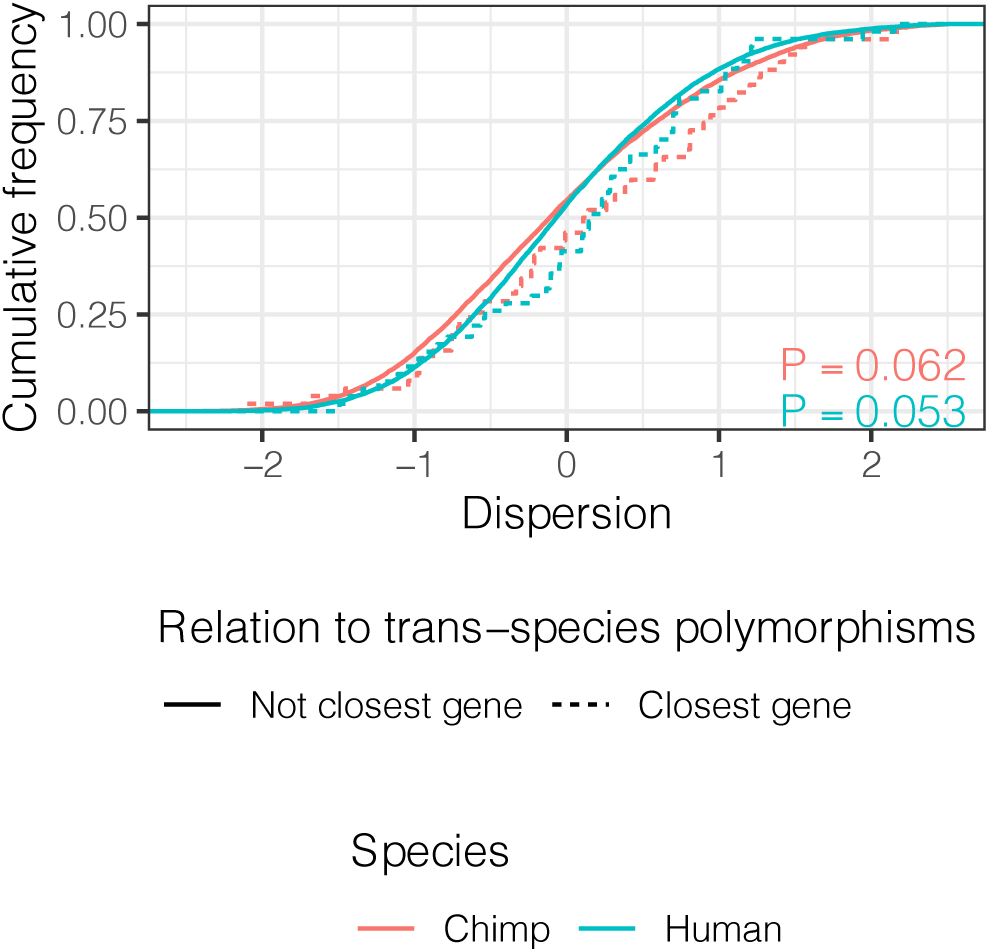
Genes closest to trans-species polymorphisms exhibit similar dispersion. ECDF of dispersion estimates in chimpanzee and human of these genes (defined as closest protein coding gene to trans-species SNP and <100kb away). P-values represent a two-sided Mann-Whitney U-test for each species.

A previous study identified 125 trans-species polymorphic haplotypes outside of the MHC region that are shared between chimpanzee and human, all but two of which are in noncoding regions (Leffler et al., 2013). Whether these polymorphisms are maintained by balancing selection because of their potential for regulatory effects on gene expression is not clear. We found that the set of genes nearest to these trans-species SNPs have higher median levels of dispersion than distal genes, though the effect is small and may be due to chance (*P*=0.053, FigS11). If these trans-species SNPs have conserved regulatory activity, which diversifies the expression levels of nearby genes, we would additionally expect to see similar eQTL effects in both human and chimpanzee. To test this, we remapped eQTLs for these SNPs in both species with a uniform pipeline (see Methods). We detected 37/192 trans-species SNPs with a clear eQTL signal (FDR<0.1) in the well-powered human dataset, such as rs257899, which associates with *SLC27A6* expression. However, we did not identify any significant *cis* eQTL activity for this SNP in chimpanzee (Fig7A). More generally, we did not find any inter-species correlation of regulatory effect size estimates among the 12 trans-species haplotypes with an eQTL in human (FDR<0.1; Fig7B) that were also tested in chimp (Methods). Considering all trans-species SNPs, we did not find any evidence for their regulatory effects in chimpanzee, compared to a set of control SNPs (Fig7C). While we found these SNPs to have measurable regulatory activity in the human dataset, it was not significantly different than that of control SNPs (Fig7D).

**Fig7.**
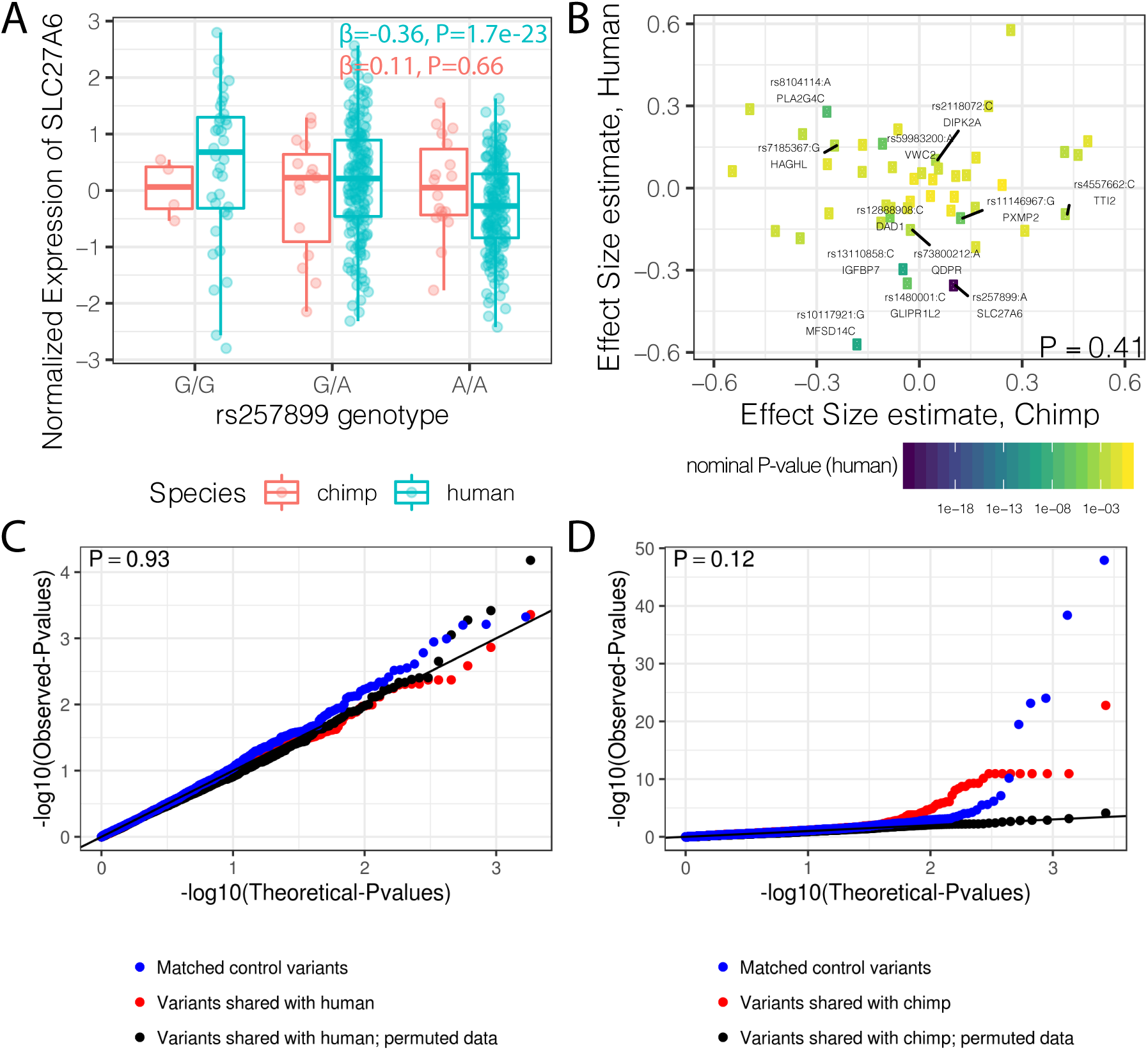
Trans-species polymorphisms do not detectably regulate gene expression. (A) Boxplot of *SLC27A6* expression stratified by species and genotype of the trans-species SNP rs257899, the most significant eQTL of the trans-species polymorphisms tested in human heart (left ventricle) in GTEx. eQTL effect size estimates (β) and nominal P-values are provided. (B) There is not a general correlation of gene regulation effects of trans-species SNPs between chimpanzee and human. For each trans-species polymorphic region previously identified (Leffler et al., 2013), the most significant SNP:gene pair in human is shown with the effect size estimate in both human and chimpanzee. Only the 48 regions where the strongest human SNP:gene association was also a one-to-one ortholog and the same SNP:gene pair was also tested in chimpanzee were plotted. Labelled SNP:gene pairs indicate FDR<0.1 in human. One sided P-value provided for Pearson correlation, under the alternative hypothesis that effect sizes should be positively correlated between species. Only SNP:gene pairs FDR<0.1 in human were considered for this test. (C) The trans-species polymorphisms do not have detectable *cis* eQTL activity in chimpanzee. QQ-plot of P-values of *cis* eQTL activity of the trans-species polymorphisms, compared to a sample permutation control, and to a control set of SNPs. (D) Same as (C) but testing *cis* eQTL activity in human. P-values provided for (C) and (D) represent one-sided Mann-Whitney U-test with the alternative hypothesis that trans-species polymorphisms have smaller *cis* eQTL P-values than the control SNPs.

Finally, we asked whether these SNPs may be under selection due to eQTL effects in tissues other than heart. To this end, we identified the most significant eQTL P-value for each of these SNPs across all GTEx tissues and found that these eQTL effects are not statistically different from that of control SNPs (FigS12A). Furthermore, the tissues where the eQTL minimum P-value was identified are similar to those of control SNPs, suggesting these trans-species SNPs in general do not have specific regulatory activity in any particular tissue. In summary, we found no compelling evidence that these trans-species polymorphisms have strong regulatory activity in either species.

**FigS12.**
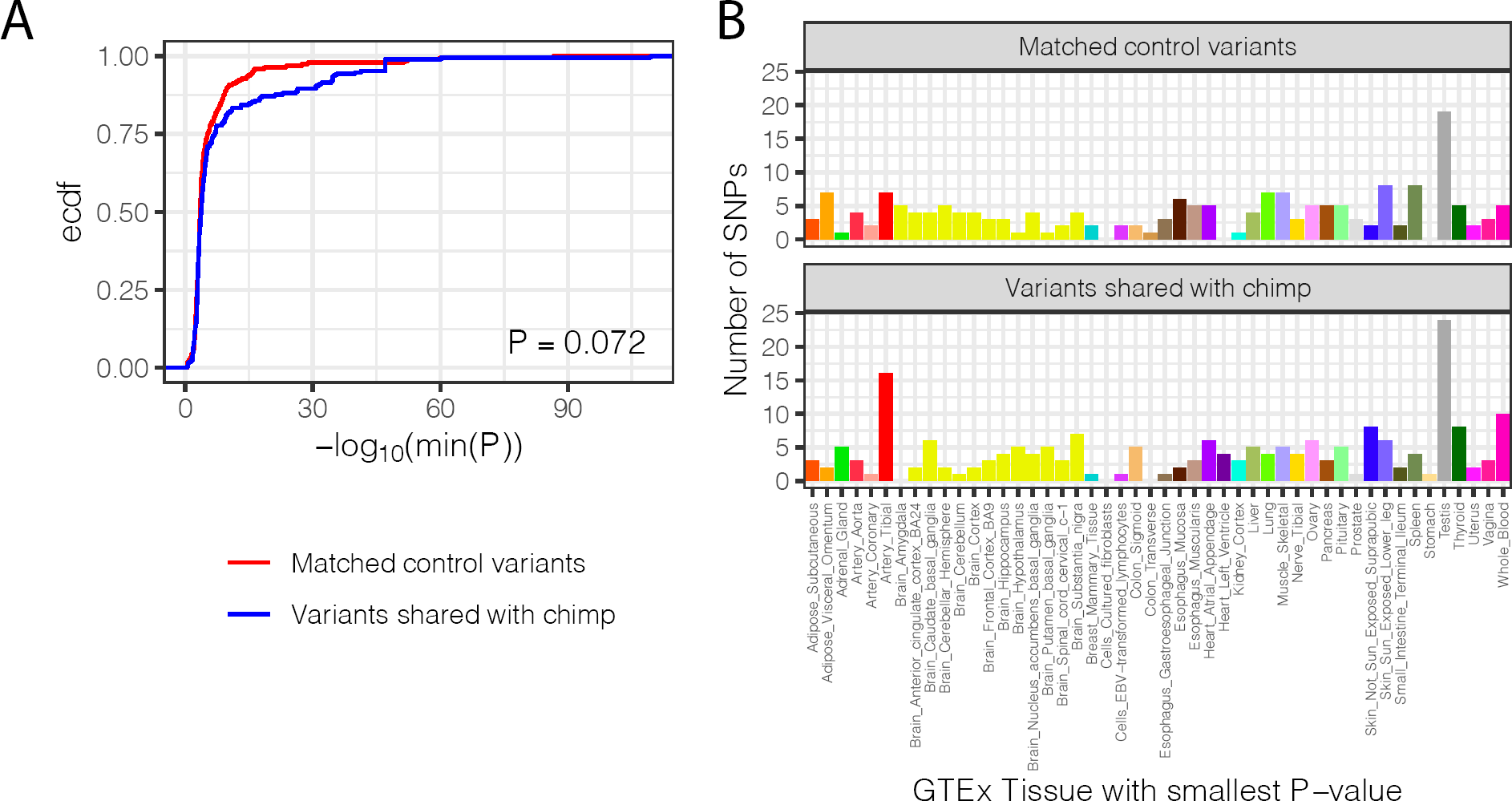
Trans-species polymorphisms do not regulate gene expression differently than control SNPs across all GTEx tissues. (A) The distribution of the smallest eQTL P-value in GTEx tissues for each trans-species polymorphism, compared to the matched control set of SNPs (<100kb from trans-species polymorphism but unlinked and matched for allele frequency; see Methods). (B) The distribution of GTEx tissues of the smallest eQTL P-value for each trans-species polymorphism, compared to matched control set of SNPs.

## Discussion

We set out to understand the properties that are associated with different levels of gene expression variability in human and chimpanzee populations. Because we know that gene expression variation in humans is often associated with genetic variation (in the form of eQTLs), we hypothesized that gene expression variability itself may be a trait under selection. We reasoned that a comparison of regulatory variation in humans and chimpanzees, and a comparative eQTL study, may provide evidence to support said hypothesis and further identify inter-species similarities and differences in the selective pressures on gene expression.

We found that inter-individual expression variability is highly correlated in humans and chimpanzees. At first glance, this seems to support the notion that regulatory variation evolves under similar selective pressures in both species. However, we were unable to exclude a technical explanation for this observation. It was difficult to disentangle the genetic and non-genetic contributions of this variability because we used primary tissue samples that include multiple cell types. We found that, across genes, cell type heterogeneity is a major driver of the degree to which gene expression varies in the population. Because orthologous genes in human and chimpanzee are expected to have similar expression patterns, this finding can potentially explain the observation of high correlation in expression dispersion in the two species. Though this technical explanation may be intuitive, the degree of the association between population variability and the cellular specificity of gene expression may have been overlooked without the use of single cell data.

Cell type heterogeneity has likely affected previous comparative studies of gene regulation that used primary tissue samples, including studies from our own lab. We and others have commented on this property of primary tissue comparisons in the past (Avila Cobos et al., 2018; Blekhman et al., 2008; Newman et al., 2015; Selewa et al., 2020), but without single cell data it was impossible to effectively assess the magnitude of this effect. Our findings further underscore the need for single cell measurements to disentangle sources of variation in bulk RNA-seq data, especially from primary tissues. Future work examining population variability should take this into account, possibly by collecting single cell data, to separate cell-type heterogeneity from heterogeneity within a particular cell type. It is important to account for cellular composition not only in comparative studies of variation in gene expression, but also in studies that focus on a single species. Indeed, cellular heterogeneity may itself have a genetic component, which will be confounded with regulatory differences within a cell type.

Notwithstanding these complications, our observations do indicate that natural selection has played a role in shaping inter-species similarities and differences in gene expression variability. Differences in cellular composition cannot explain the observed correlation between dispersion and measures of nucleotide divergence and diversity. This was the first observation in our study that provided some measure of support for the hypothesis that regulatory variation may be a genetic trait. Though the inference of selection at the genomic level indicates selection on coding regions (not gene regulation), the correlation with the degree of variation in expression suggests that the regulation of functionally important genes is also a selected trait.

Encouraged by this finding, we were able to find more evidence to support our hypothesis by carrying out a comparative eQTL analysis. We identified eQTLs in humans by using the GTEx data, which sampled hundreds of individuals. In chimpanzee, we identified eQTLs by using our 39 samples. It is quite difficult to obtain chimpanzee primary tissue samples and though this sample size is modest, it is probably the largest collection of chimpanzee primary tissue samples ever reported. The difference in sample size between the human and chimpanzee eQTL discovery panels means that observations of human-specific eGenes are quite expected and can often be explained by the fact that we have more power to detect eQTLs in humans. In contrast, the observation of chimpanzee-specific eGenes is quite meaningful because it is much less likely that the human sample was underpowered to detect eQTLs for said genes if they existed.

With that in mind, our observation that species-specific eGenes have higher variability in the species in which the eQTL was detected is significant, because it directly points to a genetic basis for differences in expression dispersion between humans and chimpanzees. This observation is also consistent with the notion that genetic variation within species contributes to the overall inter-species divergence in gene regulation. This is a critical piece of evidence supporting our hypothesis, though we acknowledge that under some complicated scenarios, one could evoke cellular composition as a potential explanation for this observation as well. We argue, however, that this would require inter-species differences in cellular composition to segregate with dozens of genotypes in just one species, and for these genotypes to appear as *cis* eQTLs for genes that have higher divergence due to cellular composition. The requirement for the genotypes to correlate with the difference in cellular composition while also being in proximity to the specific set of eGenes that would satisfy the divergence requirement is extremely unlikely, though we cannot offer empirical data to entirely exclude this possibility. That said, we also observed stronger signals of coding selection for non-eGenes than eGenes in both species, further suggesting selection on the eGenes themselves (this is not expected if our observations are to be explained by differences in cellular composition).

Collectively, our observations suggest that, across species, eQTLs may be a subtle indicator of dosage insensitivity for relatively neutrally evolving genes. Thus, we believe that an inter-species analysis of population variability in gene expression may be a relatively simple way to complement existing methods to assess differences in selection pressures between lineages. Interestingly, the eGenes we identified only in chimpanzee, but not in human, are slightly enriched for transcription regulatory processes, reminiscent of a previous observation that transcription factors seem to be enriched among the genes positively selected for in the human lineage (Blekhman et al., 2008; Gilad et al., 2006).

In both human and chimpanzee, we found that genes involved in immune response are among the most variably expressed and strongly enriched among species-shared eGenes. This is consistent with a body of literature (Croze et al., 2016; Ejsmond and Radwan, 2015; Hagai et al., 2018; Shultz and Sackton, 2019) that points to diversifying selective pressure on immune related cell-surface receptor genes to identify and combat diverse and ever-evolving pathogens. A prime example of this is the abundance of MHC complex genes among the most variable genes in both species, and strongly enriched in the subset of shared eGenes. However, the extent to which quantitative regulation of gene expression is functionally important to pathogen defense and thus the target of selection is unclear. Alternatively, these regulatory variants may be hitchhiking with functionally important and tightly linked coding variants under strong positive or balancing selection (D’Antonio et al., 2019; Meyer et al., 2018; Shiina et al., 2009). If non-coding trans-species polymorphisms are targets of long lived balancing selection on gene regulation (Johnsen et al., 2008; Leffler et al., 2013), we expect these polymorphisms to display similar eQTL effects in both species. However, we failed to identify generally conserved regulatory effects of trans-species polymorphisms. Though we found clear instances of regulatory effects from some of these trans-species SNPs in humans, we note that this human dataset is well powered enough to detect similar regulatory effects even from random control SNPs. The cumulative regulatory effects of trans-species polymorphisms are not significantly different than the control SNPs. We acknowledge that we may be underpowered to detect subtle conserved regulatory effects in chimpanzee. Moreover, some of these trans-species polymorphisms may have important regulatory functions; albeit this function may also be tissue-, cell-type-, or context-dependent and not present in our assessment of heart and other GTEx bulk tissues.

In summary, we performed a comparative assessment of expression variability and eQTL mapping and found signatures of stabilizing selection on gene regulaton in both species. A deeper understanding of differences in selection on gene expression may be gained by further assessing mean differences, variability, and eQTL contributions in various tissue types across primate groups. Such studies may benefit from single cell techniques, as we find strong contributions of cell-type heterogeneity in our analysis of variability, which may be biological or technical in nature.

## Methods

### Novel data generation

In total, 39 post-mortem heart tissue biopsies were collected from captive born chimpanzees, 18 of which have been previously described (Pavlovic et al., 2018). A partial pedigree suggests at least 8 first degree relationships among these individuals. Other metadata, including sex, age at death, and primate research center source of tissue, are detailed in TableS1. DNA and RNA were extracted from frozen tissues using kits (QIAGEN Cat No. 74104) or Trizol extraction. RNA-seq and whole genome sequencing libraries were prepared according to manufacturer’s protocols (PolyACapture followed by TruSeq v2 RNA Library prep kit; same RNA-seq protocol used by GTEx consortium. Nextera DNA Flex Library prep kit). Sequencing of RNA-seq libraries was performed by University of Chicago sequencing facilities on HiSeq 4000 using 75bp single end sequencing chemistry. The 10 RNA-seq libraries previously described (Pavlovic et al., 2018) were re-sequenced for additional depth, along with the 29 new libraries. Whole genome sequencing for all 39 chimpanzee samples was performed on NovaSeq using 300+300 paired end sequencing chemistry.

### RNA-seq and differential expression power analysis

39 RNA-seq fastq files for human left ventricle were chosen at random from GTEx v7 (TableS1, see acknowledgements). Additionally, the 10 human and 18 chimpanzee RNA-seq libraries previously generated (Pavlovic et al., 2018), and all the novel chimpanzee RNA-seq libraries generated in this study are described in Table S1. Fastq files for samples that were sequenced on multiple lanes were combined after confirmation that all gene expression profiles for all fastq files cluster primarily by sample and not by sequencing lane. GTEx-derived fastq files were trimmed to 75bp single-end reads to match the non-GTEx sequencing data. Reads were aligned to the appropriate annotated genome (GRCh38.p13 or Pan_tro_3.0 from Ensembl release 95 (Zerbino et al., 2018)) using STAR aligner (Dobin et al., 2013) default parameters. Only chromosomal contigs were considered for read alignment throughout this work. That is, unplaced contigs were excluded from the reference genome. Gene counts were obtained with subread featureCounts (Liao et al., 2014) using a previously described annotation file of human-chimpanzee orthologous exons (Pavlovic et al., 2018). The gene expression matrix was converted to CountsPerMillion (CPM) with edgeR (Robinson et al., 2010) to normalize reads to library size. The mean-variance trend was estimated using limma-voom (Ritchie et al., 2015). Genes with less than 6 CPM in all samples were excluded from further analysis. This cutoff was chosen based on visual inspection of the voom mean-variance trend to identify where the trend becomes unstable. To normalize differences in orthologous exonic gene size between human and chimpanzee, we converted the log(CPM) matrix to log(RPKM) based on the species-specific length of orthologous exonic regions. We then visually inspected PCA plots and hierarchical clustering to identify potential outliers and batch effects (FigS1). We note that although all the chimpanzee RNA isolation and sequencing were prepared separately from GTEx heart samples, PCA and clustering analysis suggest that the inter-species differences vastly outweigh the technical batch effects (based on the inter-species but within-batch samples sourced from Pavlovic et al., 2018). The dataset of 49 human samples and 39 chimpanzee samples was culled to 39 human and 39 chimpanzee samples based on exclusion of 5 obvious human outliers which did not cluster with the rest and had among the lowest read depths (FigS1A, FigS1B). The remaining human samples to exclude to reach a balanced set of 39 human and 39 chimpanzee samples were chosen by excluding the remaining GTEx samples with the lowest mapped read depths. Differential expression was tested using limma (Ritchie et al., 2015), using the eBayes function with default parameters and applying Benjamini-Hochberg FDR estimation. For FigS3, this process was repeated at varying sample size (sampling with replacement) and at various read depths. Sequencing depth subsampling analysis was performed at the level of bam files to obtain matched numbers of mapped reads across samples and differential expression analyses was repeated.

### Expression variability estimation

To estimate the mean and variance of gene expression, we assume

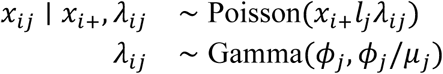

where *x*_*ij*_ is the number of reads mapping to gene *j* in sample *i* (*i* = 1, …, *n*; *j* = 1, …, *p*), *x*_*i*+_ = ∑_*j*_ *x*_*ij*_ is the total number of reads observed in sample *i, l*_*j*_ is the effective length (Pachter, 2011) of gene *j*, and *λ*_*ij*_ is the true relative gene expression of gene *j* in sample *i*. The effective length for each gene, *l*_*j*_, was calculated separately for chimpanzee and human as the length of orthologous exonic regions from which aligned reads were summed to create a count matrix. Under this model, true gene expression values for gene *j, λ*_1*j*_, …, *λ*_*nj*_, follow a Gamma distribution with mean *μ*_*j*_ and variance 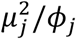, implying that the observed counts *x*_*ij*_, …, *x*_*nj*_ follow a Negative Binomial distribution with mean *x*_*i*+_*l*_*j*_*μ*_*j*_ and overdispersion 1/*ϕ*_*j*_. This model corresponds to a generalized linear model (Hilbe, 2014), which we fit by maximizing the likelihood using the ‘glm.nb’ function in the R package *MASS*.

We fit a LOESS trend using the ‘loess.fit’ function in R (with degree=1) to the mean-overdispersion trend across all genes and considered the residual from the trend as the gene’s mean-corrected dispersion. We estimated standard errors for each dispersion parameter by bootstrapping with replacement 1000 times. We estimated bootstrap p-values to test the alternative hypothesis that the absolute difference in dispersion between chimpanzee and human is greater than zero, bootstrapping with replacement 10,000 times. More specifically, we estimated the distribution of the absolute difference in dispersion under the null by performing 10,000 iterations of resamples (n=39 individuals) from a joint count matrix containing all human and chimpanzee individuals. P-values for each gene are then defined as the fraction of resamples with an absolute difference greater than what is observed. False discovery rates were estimated using Storey’s q-value (Storey and Tibshirani, 2003).

### Dispersion and cell-type heterogeneity

Single cell RNA-seq data were downloaded from the *Tabula Muris* mouse single cell atlas (The Tabula Muris Consortium et al., 2018). This dataset contains both FACS-based and droplet-based single cell RNA-seq datasets for adult mouse heart. Only the FACS based heart data were used, as the droplet based data are much sparser by comparison (The Tabula Muris Consortium et al., 2018). Data were analyzed with Seurat (Butler et al., 2018) using the published cell type labels (The Tabula Muris Consortium et al., 2018). After subsetting cells that contain at least 1000 genes with nonzero counts, the scTransform function was used to obtain a normalized count matrix used for plotting Fig3A-B. A cell-type-specificity score, τ (Kryuchkova-Mostacci and Robinson-Rechavi, 2016), was calculated for each gene (only considering one-to-one mouse/human orthologs) by utilizing the 9 cell type labels assigned by *Tabula Muris* to sum raw read counts from each cell type to create a pseudo-bulk count matrix. The pseudo-bulk count matrix was converted to CountsPerMillion and subsequently used to calculate τ with open source software (doi:10.5281/zenodo.3558708). CIBERSORT (Newman et al., 2015) was used to estimate cell types in the bulk RNA-seq datasets used for the RNA-seq dispersion and power analyses. The top 200 marker genes (Seurat FindMarkers function) unique to each cell type were used as signature genes for CIBERSORT.

### Chimpanzee genome sequencing

For whole genome sequencing data processing, we followed general guidelines for read alignment and *de novo* variant calling as previously described (Li, 2014). More precise steps are as follows: Sequencing adapters were trimmed with cutadapt (Martin, 2011) and aligned to Pan_tro_3.0 (Ensembl) using bwa-aligner (Li and Durbin, 2009) with default parameters. The average genome coverage in every sample was >30X. Sample-specific statistics for basic data processing steps, including read alignment and variant calling, are summarized in tableS7. PCR duplicates were removed via Picard tools. Low-complexity regions of the genome were determined using dustmasker (Morgulis et al., 2006) with default settings and excluded from variant calling. Sites where any sample had coverage (after PCR duplicate removal) outside of d+3√d and d+4√d coverage (where d is the sample average fold coverage across the genome) were also excluded, as these regions are enriched for duplicated or paralogous regions which are prone to false heterozygous variant calls (Li, 2014). The resulting callable sites span 2,544,417,587 out of 2,967,125,077 bases on the contiguous chromosomal genome. Variants were called in all samples jointly using freebayes (Garrison and Marth, 2012) with the following parameters: {--min-coverage 3 --max-coverage 150 -k –standard-filters -n 2 –report-genotype-likelihood-max} and subsequently filtered for Phred-scaled quality score > 30 to generate a VCF file (see Data Availability). Due to memory constraints, variant calling was executed in 2.5 megabase chunks which were later merged. In total, 19,789,407 single nucleotide variants passed variant calling filters, yielding a transition/transversion ratio of 2.08. Variants in the accompanying VCF and throughout this work were left ‘clumped, meaning that completely linked SNPs within 3bp (the default setting of freebayes) are combined into a single variant in the VCF, and tested as a single variant during eQTL calling.

For admixture analysis, a VCF file of previously sequenced wild-born chimpanzees (de Manuel et al., 2016) was lifted over to Pan_tro_3.0 and merged with the VCF file described above, keeping only variants present in both sets. Variants in high LD were pruned using plink (Purcell et al., 2007) with parameters {--indep-pairwise 50 5 0.5}. The resulting genotypes were analyzed for population structure with PCA using ‘prcomp’ function in R. Additionally, we utilized Admixture software (Alexander et al., 2009) with K=4 clusters, as there are 4 recognized distinct chimpanzee subspecies, all of which were represented in the wild-born cohort (de Manuel et al., 2016).

### Cis eQTLmapping (chimpanzee)

RNA-seq reads were re-aligned with STAR aligner (Dobin et al., 2013) using Pan_tro_3.0 gene annotations from Ensembl version 95 (Zerbino et al., 2018). Gene counts were quantified with STAR --quantMode GeneCounts to compile a gene expression matrix. Genes were filtered to require at least 8 read counts in 75% of samples, leaving 13,545 genes for further analysis. The resulting read count matrix was converted to log(CPM), standardized across individuals, and quantile normalized to a normal distribution across genes as previously described (Degner et al., 2012). Principle component analysis of the normalized matrix identified significant associations between various observed technical factors and some of the first 10 principle components, including RNA library prep batch and sex, and as such, principle components were included later as covariates during eQTL calling.

Variants were filtered for MAF>0.1 and Hardy-Weinberg equilibrium (nominal P>1×10^−7.5^, hardy function in plink) to filter out rare variants and genotyping errors. The resulting 5,957,179 variants were each tested for association with expression of each local *cis* gene (*cis* window defined as within 250kb of gene body). MatrixEQTL (Shabalin, 2012) was used to implement a linear mixed model for each *cis* variant:gene pair to estimate the effect of the variant genotype on normalized expression gene expression. We supplied MatrixEQTL with a genetic relatedness matrix (GRM) to account for heteroskedastic errors generated from underlying population structure and genetic relatedness amongst individuals. The standardized GRM was produced by GEMMA (Zhou and Stephens, 2012) using variants pruned for LD as described for admixture analysis. Additionally, between 0 and 15 gene expression principle components (PCs) were tested as covariates to the linear model and 10 PCs were included in the final model as this maximizes the number of eQTLs (FigS5A). Manual inspection of normalized expression boxplots stratified by genotype for the top eQTLs revealed many of the nominally strongest associations were driven by a single expression outlier point for a single homozygous individual, MD_And. Further inspection revealed this individual has among the highest levels of homozygosity genome-wide among our chimpanzee cohort (TableS7), possibly due to inbreeding. Given that this individual is the only chimpanzee sourced from MD Anderson primate research center, we felt justified excluding this individual from eQTL calling to minimize false associations. After excluding this individual and re-performing eQTL mapping (n=38), visual inspection of a QQ-plot of P-values compared to permuted null data (where the sample labels for expression and covariates were randomly assigned to genotype) indicates inflation of small P-values, and that P-values are well calibrated under the permuted null (FigS5B). To obtain gene-level P-values that test whether a gene contains an eQTL (eGenes) we used EigenMT, a method which approximates permutation testing procedures to account for multiple testing of linked SNPs (Davis et al., 2016).

### Cis eQTL mapping (human)

We downloaded gene-level (eGene) summary statistics for Heart_Left_ventricle GTEx v8 from GTEx portal (https://gtexportal.org/). The summary statistics from this mapping pipeline only considers expressed genes, which are defined as >0.1 TPM in at least 20% of samples and ≥6 reads in at least 20% of samples. The analysis in FigS9B required more than summary statistics. We downloaded normalized phenotypes, covariates, and genotypes for GTEx v8 data for Heart_Left_ventricle and remapped eGenes using varying sample sizes using a mapping pipeline nearly identical to GTEx. Namely, we used FastQTL (Ongen et al., 2016) on the supplied data with randomly selected individuals corresponding to sample sizes (n) of 40, 60, 80, 100, 120, 160, 200. Similar to guidelines described by GTEx (https://gtexportal.org/), we included only 10 PEER factor covariates for n=40, 15 for 40>n>150, or 30 for n>150.

### Gene-wise conservation statistics

Gene-wise amino acid percent identity between chimpanzee and human was obtained from BioMart (Kinsella et al., 2011). Gene-wise pan-mammal dn/ds for each gene was obtained from a previous study that used alignments from 29 mammals (Broad Institute Sequencing Platform and Whole Genome Assembly Team et al., 2011). P_n_/P_s_ was calculated from all GTEx v8 genotype data and the union of all *Pan troglodyte* genotypes available in this study and DeManuel et al (de Manuel et al., 2016). Specifically, Ensembl vep (McLaren et al., 2016) was used to annotate coding variants with MAF>0.1 as synonymous or non-synonymous to tabulate Pn (number of polymorphic non-synonymous sites) and Ps (number of polymorphic synonymous sites) for each gene within each species. After requiring that genes have at least one polymorphism in both species, a pseudocount of 0.5 was added to both Pn and Ps for each gene for both species to avoid division-by-zero errors. TATA box genes were classified as genes with a TATA motif within 35bp of a transcription initiation site from published transcription initiation sites (Abugessaisa et al., 2019).

### eQTL-mapping of shared polymorphisms

263 SNPs among 125 regions that are trans-species polymorphisms between chimpanzee and humans were obtained from Leffler et al (Leffler et al., 2013). Each trans-species polymorphic *region* contains at least two trans-species SNPs to ensure regions are identical by descent rather than recurrent mutation of an isolated SNP (Leffler et al., 2013). Only SNPs with MAF>0.1 in our datasets were further utilized, leaving 192 SNPs in chimpanzee, 196 SNPs in human (GTEx), and 144 SNPs (amongst 76 regions) tested for eQTL activity in both species. For each test SNP, a matched control SNP was randomly chosen for each species with the criteria that it should have a matching allele frequency (±5% MAF), within 100kb of the test SNP, and unlinked to the test SNP (R^2^<0.2, LD calculated with plink). We used MatrixEQTL to retest these SNPs for *cis* eQTL activity (1MB window) and *trans* eQTL activity using MatrixEQTL with the same normalized gene expression matrix, GRM matrix (for chimpanzee only), and covariates described above for chimpanzee and human *cis* eQTL mapping. The effect sizes between species occasionally had to be re-polarized to relate to the same allele, as the effect sizes obtained from eQTL testing software is often polarized by minor allele or by reference vs non-reference allele, though the minor allele and/or reference allele at these trans-species polymorphisms is not always the same between species.

### Gene set enrichment analysis

Gene set enrichment and gene ontology overlap analysis was performed with clusterProfiler R package (Yu et al., 2012). The GSEA test was performed with an ordered gene list using the ‘gseGO’ function with 1,000,000 permutations. As ordering genes was not applicable to inter-species eGene classifications, the ‘enrichGO’ function was used to perform gene ontology overlap analysis (hypergeometric test) with foreground and background gene sets based on eGene classifications.

## Supplemental Tables

TableS1: RNA-seq datasets used, with accession numbers for new and published datasets, number and percent of reads mapped, and some other basic stats

TableS2: DE analysis results with full 39 human + 39 chimpanzee dataset. Per gene effect sizes, P-values, etc

TableS3: expression and dispersion estimates for chimpanzee and human

TableS4: GSEA of high vs low dispersion genes based on human

TableS5: GSEA of human-chimpanzee difference in variability

TableS6: GSEA of human-chimpanzee difference in variability when 5 HBV/HCV chimpanzees excluded from dispersion estimates

TableS7: Per sample summary stats for genome sequencing, including coverage, heterozygosity, tissue source, chimpanzee IDs, etc

TableS8: Summary stats for chimpanzee eGenes

TableS9: Gene ontology enrichment for shared eGenes

TableS10: Gene ontology enrichment for human specific eGenes

## Acknowledgements

We thank Natalia Gonzales, Michelle Ward and other members of the Gilad lab for helpful discussions and comments on the manuscript. This work was supported by NIH grant R35GM131726 as well as the Yerkes National Primate Research Center Base Grant ORIP/OD P51OD011132. Computational resources were provided by the University of Chicago Research Computing Center

The Genotype-Tissue Expression (GTEx) Project was supported by the <u>Common Fund</u> of the Office of the Director of the National Institutes of Health, and by NCI, NHGRI, NHLBI, NIDA, NIMH, and NINDS. The data used for the analyses described in this manuscript were obtained from: the GTEx Portal on 10/08/19 and/or dbGaP accession number phs000424.v7.p2 on 09/17/19.

## Data Availability

Novel RNA-seq data is made available under Gene Expression Omnibus (GEO) accession number GSE151397. Chimpanzee whole genome sequencing data is available under Sequence Read Archive (SRA) accession number PRJNA635393 (raw data) and European Variation Archive (EVA) accession number XXX (variant calls). Reproducible code available at https://github.com/bfairkun/Comparative_eQTL.

## References

Abugessaisa I, Noguchi S, Hasegawa A, Kondo A, Kawaji H, Carninci P, Kasukawa T. 2019. refTSS: A Reference Data Set for Human and Mouse Transcription Start Sites. Journal of Molecular Biology 431:2407–2422. doi:10.1016/j.jmb.2019.04.045

Albert FW, Bloom JS, Siegel J, Day L, Kruglyak L. 2018. Genetics of trans-regulatory variation in gene expression. eLife 7:e35471. doi:10.7554/eLife.35471

Alexander DH, Novembre J, Lange K. 2009. Fast model-based estimation of ancestry in unrelated individuals. Genome Research 19:1655–1664. doi:10.1101/gr.094052.109

Avila Cobos F, Vandesompele J, Mestdagh P, De Preter K. 2018. Computational deconvolution of transcriptomics data from mixed cell populations. Bioinformatics 34:1969–1979. doi:10.1093/bioinformatics/bty019

Barbosa-Morais NL, Irimia M, Pan Q, Xiong HY, Gueroussov S, Lee LJ, Slobodeniuc V, Kutter C, Watt S, Colak R, Kim T, Misquitta-Ali CM, Wilson MD, Kim PM, Odom DT, Frey BJ, Blencowe BJ. 2012. The Evolutionary Landscape of Alternative Splicing in Vertebrate Species. Science 338:1587–1593. doi:10.1126/science.1230612

Bashkeel N, Perkins TJ, Kærn M, Lee JM. 2019. Human gene expression variability and its dependence on methylation and aging. BMC Genomics 20:941. doi:10.1186/s12864-019-6308-7

Battle A, Mostafavi S, Zhu X, Potash JB, Weissman MM, McCormick C, Haudenschild CD, Beckman KB, Shi J, Mei R, Urban AE, Montgomery SB, Levinson DF, Koller D. 2014. Characterizing the genetic basis of transcriptome diversity through RNA-sequencing of 922 individuals. Genome Research 24:14–24. doi:10.1101/gr.155192.113

Blake WJ, Balázsi G, Kohanski MA, Isaacs FJ, Murphy KF, Kuang Y, Cantor CR, Walt DR, Collins JJ. 2006. Phenotypic Consequences of Promoter-Mediated Transcriptional Noise. Molecular Cell 24:853–865. doi:10.1016/j.molcel.2006.11.003

Blekhman R, Oshlack A, Chabot AE, Smyth GK, Gilad Y. 2008. Gene Regulation in Primates Evolves under Tissue-Specific Selection Pressures. PLoS Genet 4:e1000271. doi:10.1371/journal.pgen.1000271

BLUEPRINT Consortium, Ecker S, Chen L, Pancaldi V, Bagger FO, Fernández JM, Carrillo de Santa Pau E, Juan D, Mann AL, Watt S, Casale FP, Sidiropoulos N, Rapin N, Merkel A, Stunnenberg HG, Stegle O, Frontini M, Downes K, Pastinen T, Kuijpers TW, Rico D, Valencia A, Beck S, Soranzo N, Paul DS. 2017. Genome-wide analysis of differential transcriptional and epigenetic variability across human immune cell types. Genome Biol 18:18. doi:10.1186/s13059-017-1156-8

Bódi Z, Farkas Z, Nevozhay D, Kalapis D, Lázár V, Csörgő B, Nyerges Á, Szamecz B, Fekete G, Papp B, Araújo H, Oliveira JL, Moura G, Santos MAS, Székely Jr T, Balázsi G, Pál C. 2017. Phenotypic heterogeneity promotes adaptive evolution. PLoS Biol 15:e2000644. doi:10.1371/journal.pbio.2000644

Brawand D, Soumillon M, Necsulea A, Julien P, Csárdi G, Harrigan P, Weier M, Liechti A, Aximu-Petri A, Kircher M, Albert FW, Zeller U, Khaitovich P, Grützner F, Bergmann S, Nielsen R, Pääbo S, Kaessmann H. 2011. The evolution of gene expression levels in mammalian organs. Nature 478:343–348. doi:10.1038/nature10532

Broad Institute Sequencing Platform and Whole Genome Assembly Team, Baylor College of Medicine Human Genome Sequencing Center Sequencing Team, Genome Institute at Washington University, Lindblad-Toh K, Garber M, Zuk O, Lin MF, Parker BJ, Washietl S, Kheradpour P, Ernst J, Jordan G, Mauceli E, Ward LD, Lowe CB, Holloway AK, Clamp M, Gnerre S, Alföldi J, Beal K, Chang J, Clawson H, Cuff J, Di Palma F, Fitzgerald S, Flicek P, Guttman M, Hubisz MJ, Jaffe DB, Jungreis I, Kent WJ, Kostka D, Lara M, Martins AL, Massingham T, Moltke I, Raney BJ, Rasmussen MD, Robinson J, Stark A, Vilella AJ, Wen J, Xie X, Zody MC, Worley KC, Kovar CL, Muzny DM, Gibbs RA, Warren WC, Mardis ER, Weinstock GM, Wilson RK, Birney E, Margulies EH, Herrero J, Green ED, Haussler D, Siepel A, Goldman N, Pollard KS, Pedersen JS, Lander ES, Kellis M. 2011. A high-resolution map of human evolutionary constraint using 29 mammals. Nature 478:476–482. doi:10.1038/nature10530

Butler A, Hoffman P, Smibert P, Papalexi E, Satija R. 2018. Integrating single-cell transcriptomic data across different conditions, technologies, and species. Nat Biotechnol 36:411–420. doi:10.1038/nbt.4096

Chan ET, Quon GT, Chua G, Babak T, Trochesset M, Zirngibl RA, Aubin J, Ratcliffe MJ, Wilde A, Brudno M, Morris QD, Hughes TR. 2009. Conservation of core gene expression in vertebrate tissues. J Biol 8:33. doi:10.1186/jbiol130

Croze M, Živković D, Stephan W, Hutter S. 2016. Balancing selection on immunity genes: review of the current literature and new analysis in Drosophila melanogaster. Zoology 119:322–329. doi:10.1016/j.zool.2016.03.004

D’Antonio M, Reyna J, Jakubosky D, Donovan MK, Bonder M-J, Matsui H, Stegle O, Nariai N, D’Antonio-Chronowska A, Frazer KA. 2019. Systematic genetic analysis of the MHC region reveals mechanistic underpinnings of HLA type associations with disease. eLife 8:e48476. doi:10.7554/eLife.48476

Davis JR, Fresard L, Knowles DA, Pala M, Bustamante CD, Battle A, Montgomery SB. 2016. An Efficient Multiple-Testing Adjustment for eQTL Studies that Accounts for Linkage Disequilibrium between Variants. The American Journal of Human Genetics 98:216–224. doi:10.1016/j.ajhg.2015.11.021

de Jong TV, Moshkin YM, Guryev V. 2019. Gene expression variability: the other dimension in transcriptome analysis. Physiological Genomics 51:145–158. doi:10.1152/physiolgenomics.00128.2018

de Manuel M, Kuhlwilm M, Frandsen P, Sousa VC, Desai T, Prado-Martinez J, Hernandez-Rodriguez J, Dupanloup I, Lao O, Hallast P, Schmidt JM, Heredia-Genestar JM, Benazzo A, Barbujani G, Peter BM, Kuderna LFK, Casals F, Angedakin S, Arandjelovic M, Boesch C, Kuhl H, Vigilant L, Langergraber K, Novembre J, Gut M, Gut I, Navarro A, Carlsen F, Andres AM, Siegismund HR, Scally A, Excoffier L, Tyler-Smith C, Castellano S, Xue Y, Hvilsom C, Marques-Bonet T. 2016. Chimpanzee genomic diversity reveals ancient admixture with bonobos. Science 354:477–481. doi:10.1126/science.aag2602

Degner JF, Pai AA, Pique-Regi R, Veyrieras J-B, Gaffney DJ, Pickrell JK, De Leon S, Michelini K, Lewellen N, Crawford GE, Stephens M, Gilad Y, Pritchard JK. 2012. DNase I sensitivity QTLs are a major determinant of human expression variation. Nature 482:390–394. doi:10.1038/nature10808

Dobin A, Davis CA, Schlesinger F, Drenkow J, Zaleski C, Jha S, Batut P, Chaisson M, Gingeras TR. 2013. STAR: ultrafast universal RNA-seq aligner. Bioinformatics 29:15–21. doi:10.1093/bioinformatics/bts635

Ejsmond MJ, Radwan J. 2015. Red Queen Processes Drive Positive Selection on Major Histocompatibility Complex (MHC) Genes. PLoS Comput Biol 11:e1004627. doi:10.1371/journal.pcbi.1004627

Eling N, Richard AC, Richardson S, Marioni JC, Vallejos CA. 2018. Correcting the Mean-Variance Dependency for Differential Variability Testing Using Single-Cell RNA Sequencing Data. Cell Systems 7:284-294.e12. doi:10.1016/j.cels.2018.06.011

Exome Aggregation Consortium, Lek M, Karczewski KJ, Minikel EV, Samocha KE, Banks E, Fennell T, O’Donnell-Luria AH, Ware JS, Hill AJ, Cummings BB, Tukiainen T, Birnbaum DP, Kosmicki JA, Duncan LE, Estrada K, Zhao F, Zou J, Pierce-Hoffman E, Berghout J, Cooper DN, Deflaux N, DePristo M, Do R, Flannick J, Fromer M, Gauthier L, Goldstein J, Gupta N, Howrigan D, Kiezun A, Kurki MI, Moonshine AL, Natarajan P, Orozco L, Peloso GM, Poplin R, Rivas MA, Ruano-Rubio V, Rose SA, Ruderfer DM, Shakir K, Stenson PD, Stevens C, Thomas BP, Tiao G, Tusie-Luna MT, Weisburd B, Won H-H, Yu D, Altshuler DM, Ardissino D, Boehnke M, Danesh J, Donnelly S, Elosua R, Florez JC, Gabriel SB, Getz G, Glatt SJ, Hultman CM, Kathiresan S, Laakso M, McCarroll S, McCarthy MI, McGovern D, McPherson R, Neale BM, Palotie A, Purcell SM, Saleheen D, Scharf JM, Sklar P, Sullivan PF, Tuomilehto J, Tsuang MT, Watkins HC, Wilson JG, Daly MJ, MacArthur DG. 2016. Analysis of protein-coding genetic variation in 60,706 humans. Nature 536:285–291. doi:10.1038/nature19057

Fuller ZL, Niño EL, Patch HM, Bedoya-Reina OC, Baumgarten T, Muli E, Mumoki F, Ratan A, McGraw J, Frazier M, Masiga D, Schuster S, Grozinger CM, Miller W. 2015. Genome-wide analysis of signatures of selection in populations of African honey bees (Apis mellifera) using new web-based tools. BMC Genomics 16:518. doi:10.1186/s12864-015-1712-0

Garrison E, Marth G. 2012. Haplotype-based variant detection from short-read sequencing. 12073907 [q-bio].

Gilad Y, Oshlack A, Smyth GK, Speed TP, White KP. 2006. Expression profiling in primates reveals a rapid evolution of human transcription factors. Nature 440:242–245. doi:10.1038/nature04559

Glassberg EC, Gao Z, Harpak A, Lan X, Pritchard JK. 2019. Evidence for Weak Selective Constraint on Human Gene Expression. Genetics 211:757–772. doi:10.1534/genetics.118.301833

Göring HHH, Terwilliger JD, Blangero J. 2001. Large Upward Bias in Estimation of Locus-Specific Effects from Genomewide Scans. The American Journal of Human Genetics 69:1357–1369. doi:10.1086/324471

Hagai T, Chen X, Miragaia RJ, Rostom R, Gomes T, Kunowska N, Henriksson J, Park J-E, Proserpio V, Donati G, Bossini-Castillo L, Vieira Braga FA, Naamati G, Fletcher J, Stephenson E, Vegh P, Trynka G, Kondova I, Dennis M, Haniffa M, Nourmohammad A, Lässig M, Teichmann SA. 2018. Gene expression variability across cells and species shapes innate immunity. Nature 563:197–202. doi:10.1038/s41586-018-0657-2

Hilbe JM. 2014. Modeling Count Data. Cambridge: Cambridge University Press. doi:10.1017/CBO9781139236065

Ho JWK, Stefani M, dos Remedios CG, Charleston MA. 2008. Differential variability analysis of gene expression and its application to human diseases. Bioinformatics 24:i390–i398. doi:10.1093/bioinformatics/btn142

Huguet G, Nava C, Lemière N, Patin E, Laval G, Ey E, Brice A, Leboyer M, Szepetowski P, Gillberg C, Depienne C, Delorme R, Bourgeron T. 2014. Heterogeneous Pattern of Selective Pressure for PRRT2 in Human Populations, but No Association with Autism Spectrum Disorders. PLoS ONE 9:e88600. doi:10.1371/journal.pone.0088600

Ioannidis JPA. 2008. Why Most Discovered True Associations Are Inflated: Epidemiology 19:640–648. doi:10.1097/EDE.0b013e31818131e7

Jasinska AJ, Zelaya I, Service SK, Peterson CB, Cantor RM, Choi O-W, DeYoung J, Eskin E, Fairbanks LA, Fears S, Furterer AE, Huang YS, Ramensky V, Schmitt CA, Svardal H, Jorgensen MJ, Kaplan JR, Villar D, Aken BL, Flicek P, Nag R, Wong ES, Blangero J, Dyer TD, Bogomolov M, Benjamini Y, Weinstock GM, Dewar K, Sabatti C, Wilson RK, Jentsch JD, Warren W, Coppola G, Woods RP, Freimer NB. 2017. Genetic variation and gene expression across multiple tissues and developmental stages in a nonhuman primate. Nat Genet 49:1714–1721. doi:10.1038/ng.3959

Johnsen JM, Teschke M, Pavlidis P, McGee BM, Tautz D, Ginsburg D, Baines JF. 2008. Selection on cis-Regulatory Variation at B4galnt2 and Its Influence on von Willebrand Factor in House Mice. Molecular Biology and Evolution 26:567–578. doi:10.1093/molbev/msn284

Kajstura J, Leri A, Finato N, Di Loreto C, Beltrami CA, Anversa P. 1998. Myocyte proliferation in end-stage cardiac failure in humans. Proceedings of the National Academy of Sciences 95:8801–8805. doi:10.1073/pnas.95.15.8801

Khaitovich P, Enard W, Lachmann M, Pääbo S. 2006. Evolution of primate gene expression. Nat Rev Genet 7:693–702. doi:10.1038/nrg1940

Khan Z, Ford MJ, Cusanovich DA, Mitrano A, Pritchard JK, Gilad Y. 2013. Primate Transcript and Protein Expression Levels Evolve Under Compensatory Selection Pressures. Science 342:1100–1104. doi:10.1126/science.1242379

Kimura W, Nakada Y, Sadek HA. 2017. Hypoxia-induced myocardial regeneration. Journal of Applied Physiology 123:1676–1681. doi:10.1152/japplphysiol.00328.2017

Kinsella RJ, Kahari A, Haider S, Zamora J, Proctor G, Spudich G, Almeida-King J, Staines D, Derwent P, Kerhornou A, Kersey P, Flicek P. 2011. Ensembl BioMarts: a hub for data retrieval across taxonomic space. Database 2011:bar030–bar030. doi:10.1093/database/bar030

Knowles DA, Burrows CK, Blischak JD, Patterson KM, Serie DJ, Norton N, Ober C, Pritchard JK, Gilad Y. 2018. Determining the genetic basis of anthracycline-cardiotoxicity by molecular response QTL mapping in induced cardiomyocytes. eLife 7:e33480. doi:10.7554/eLife.33480

Kryuchkova-Mostacci N, Robinson-Rechavi M. 2016. A benchmark of gene expression tissue-specificity metrics. Brief Bioinform bbw008. doi:10.1093/bib/bbw008

Lam TH, Shen M, Tay MZ, Ren EC. 2017. Unique Allelic eQTL Clusters in Human MHC Haplotypes. G3 7:2595–2604. doi:10.1534/g3.117.043828

Leffler EM, Gao Z, Pfeifer S, Segurel L, Auton A, Venn O, Bowden R, Bontrop R, Wall JD, Sella G, Donnelly P, McVean G, Przeworski M. 2013. Multiple Instances of Ancient Balancing Selection Shared Between Humans and Chimpanzees. Science 339:1578–1582. doi:10.1126/science.1234070

Lemos B, Meiklejohn CD, Cáceres M, Hartl DL. 2005. Rates of divergence in gene expression profiles of primates, mice, and flies: stabilizing selection and variability among functional categories. Evolution 59:126–137.

Li H. 2014. Toward better understanding of artifacts in variant calling from high-coverage samples. Bioinformatics 30:2843–2851. doi:10.1093/bioinformatics/btu356

Li H, Durbin R. 2009. Fast and accurate short read alignment with Burrows-Wheeler transform. Bioinformatics 25:1754–1760. doi:10.1093/bioinformatics/btp324

Li J, Liu Y, Kim T, Min R, Zhang Z. 2010. Gene Expression Variability within and between Human Populations and Implications toward Disease Susceptibility. PLoS Comput Biol 6:e1000910. doi:10.1371/journal.pcbi.1000910

Liao Y, Smyth GK, Shi W. 2014. featureCounts: an efficient general purpose program for assigning sequence reads to genomic features. Bioinformatics 30:923–930. doi:10.1093/bioinformatics/btt656

Mähler N, Wang J, Terebieniec BK, Ingvarsson PK, Street NR, Hvidsten TR. 2017. Gene co-expression network connectivity is an important determinant of selective constraint. PLoS Genet 13:e1006402. doi:10.1371/journal.pgen.1006402

Marioni JC, Mason CE, Mane SM, Stephens M, Gilad Y. 2008. RNA-seq: An assessment of technical reproducibility and comparison with gene expression arrays. Genome Research 18:1509–1517. doi:10.1101/gr.079558.108

Martin M. 2011. Cutadapt removes adapter sequences from high-throughput sequencing reads. EMBnet j 17:10. doi:10.14806/ej.17.1.200

McLaren W, Gil L, Hunt SE, Riat HS, Ritchie GRS, Thormann A, Flicek P, Cunningham F. 2016. The Ensembl Variant Effect Predictor. Genome Biol 17:122. doi:10.1186/s13059-016-0974-4

Merkin J, Russell C, Chen P, Burge CB. 2012. Evolutionary dynamics of gene and isoform regulation in Mammalian tissues. Science (New York, NY) 338:1593–9. doi:10.1126/science.1228186

Meyer D, C Aguiar VR, Bitarello BD, C Brandt DY, Nunes K. 2018. A genomic perspective on HLA evolution. Immunogenetics 70:5–27. doi:10.1007/s00251-017-1017-3

Morgulis A, Gertz EM, Schäffer AA, Agarwala R. 2006. A Fast and Symmetric DUST Implementation to Mask Low-Complexity DNA Sequences. Journal of Computational Biology 13:1028–1040. doi:10.1089/cmb.2006.13.1028

Nakada Y, Canseco DC, Thet S, Abdisalaam S, Asaithamby A, Santos CX, Shah AM, Zhang H, Faber JE, Kinter MT, Szweda LI, Xing C, Hu Z, Deberardinis RJ, Schiattarella G, Hill JA, Oz O, Lu Z, Zhang CC, Kimura W, Sadek HA. 2017. Hypoxia induces heart regeneration in adult mice. Nature 541:222–227. doi:10.1038/nature20173

Newman AM, Liu CL, Green MR, Gentles AJ, Feng W, Xu Y, Hoang CD, Diehn M, Alizadeh AA. 2015. Robust enumeration of cell subsets from tissue expression profiles. Nat Methods 12:453–457. doi:10.1038/nmeth.3337

Ongen H, Buil A, Brown AA, Dermitzakis ET, Delaneau O. 2016. Fast and efficient QTL mapper for thousands of molecular phenotypes. Bioinformatics 32:1479–1485. doi:10.1093/bioinformatics/btv722

Pachter L. 2011. Models for transcript quantification from RNA-Seq. 11043889 [q-bio, stat].

Pavlovic BJ, Blake LE, Roux J, Chavarria C, Gilad Y. 2018. A Comparative Assessment of Human and Chimpanzee iPSC-derived Cardiomyocytes with Primary Heart Tissues. Sci Rep 8:15312. doi:10.1038/s41598-018-33478-9

Perry GH, Melsted P, Marioni JC, Wang Y, Bainer R, Pickrell JK, Michelini K, Zehr S, Yoder AD, Stephens M, Pritchard JK, Gilad Y. 2012. Comparative RNA sequencing reveals substantial genetic variation in endangered primates. Genome Research 22:602–610. doi:10.1101/gr.130468.111

Pickrell JK, Marioni JC, Pai AA, Degner JF, Engelhardt BE, Nkadori E, Veyrieras J-B, Stephens M, Gilad Y, Pritchard JK. 2010. Understanding mechanisms underlying human gene expression variation with RNA sequencing. Nature 464:768–772. doi:10.1038/nature08872

Popadin KY, Gutierrez-Arcelus M, Lappalainen T, Buil A, Steinberg J, Nikolaev SI, Lukowski SW, Bazykin GA, Seplyarskiy VB, Ioannidis P, Zdobnov EM, Dermitzakis ET, Antonarakis SE. 2014. Gene Age Predicts the Strength of Purifying Selection Acting on Gene Expression Variation in Humans. The American Journal of Human Genetics 95:660–674. doi:10.1016/j.ajhg.2014.11.003

Price AL, Helgason A, Thorleifsson G, McCarroll SA, Kong A, Stefansson K. 2011. Single-Tissue and Cross-Tissue Heritability of Gene Expression Via Identity-by-Descent in Related or Unrelated Individuals. PLoS Genet 7:e1001317. doi:10.1371/journal.pgen.1001317

Purcell S, Neale B, Todd-Brown K, Thomas L, Ferreira MAR, Bender D, Maller J, Sklar P, de Bakker PIW, Daly MJ, Sham PC. 2007. PLINK: A Tool Set for Whole-Genome Association and Population-Based Linkage Analyses. The American Journal of Human Genetics 81:559–575. doi:10.1086/519795

Raser JM. 2004. Control of Stochasticity in Eukaryotic Gene Expression. Science 304:1811–1814. doi:10.1126/science.1098641

Ravarani CNJ, Chalancon G, Breker M, de Groot NS, Babu MM. 2016. Affinity and competition for TBP are molecular determinants of gene expression noise. Nat Commun 7:10417. doi:10.1038/ncomms10417

Ritchie ME, Phipson B, Wu D, Hu Y, Law CW, Shi W, Smyth GK. 2015. limma powers differential expression analyses for RNA-sequencing and microarray studies. Nucleic Acids Research 43:e47–e47. doi:10.1093/nar/gkv007

Robinson MD, McCarthy DJ, Smyth GK. 2010. edgeR: a Bioconductor package for differential expression analysis of digital gene expression data. Bioinformatics 26:139–140. doi:10.1093/bioinformatics/btp616

Romero IG, Ruvinsky I, Gilad Y. 2012. Comparative studies of gene expression and the evolution of gene regulation. Nat Rev Genet 13:505–516. doi:10.1038/nrg3229

Schug J, Schuller W-P, Kappen C, Salbaum JM, Bucan M, Stoeckert CJ. 2005. Promoter features related to tissue specificity as measured by Shannon entropy. Genome Biol 6:R33. doi:10.1186/gb-2005-6-4-r33

Selewa A, Dohn R, Eckart H, Lozano S, Xie B, Gauchat E, Elorbany R, Rhodes K, Burnett J, Gilad Y, Pott S, Basu A. 2020. Systematic Comparison of High-throughput Single-Cell and Single-Nucleus Transcriptomes during Cardiomyocyte Differentiation. Sci Rep 10:1535. doi:10.1038/s41598-020-58327-6

Shabalin AA. 2012. Matrix eQTL: ultra fast eQTL analysis via large matrix operations. Bioinformatics 28:1353–1358. doi:10.1093/bioinformatics/bts163

Shiina T, Hosomichi K, Inoko H, Kulski JK. 2009. The HLA genomic loci map: expression, interaction, diversity and disease. J Hum Genet 54:15–39. doi:10.1038/jhg.2008.5

Shultz AJ, Sackton TB. 2019. Immune genes are hotspots of shared positive selection across birds and mammals. eLife 8:e41815. doi:10.7554/eLife.41815

Simonovsky E, Schuster R, Yeger-Lotem E. 2019. Large-scale analysis of human gene expression variability associates highly variable drug targets with lower drug effectiveness and safety. Bioinformatics 35:3028–3037. doi:10.1093/bioinformatics/btz023

Storey JD, Tibshirani R. 2003. Statistical significance for genomewide studies. Proceedings of the National Academy of Sciences 100:9440–9445. doi:10.1073/pnas.1530509100

Subramanian A, Tamayo P, Mootha VK, Mukherjee S, Ebert BL, Gillette MA, Paulovich A, Pomeroy SL, Golub TR, Lander ES, Mesirov JP. 2005. Gene set enrichment analysis: A knowledge-based approach for interpreting genome-wide expression profiles. Proceedings of the National Academy of Sciences 102:15545–15550. doi:10.1073/pnas.0506580102

Tanaka T, Nei M. 1989. Positive darwinian selection observed at the variable-region genes of immunoglobulins. Molecular Biology and Evolution. doi:10.1093/oxfordjournals.molbev.a040569

Těšický M, Vinkler M. 2015. Trans-Species Polymorphism in Immune Genes: General Pattern or MHC-Restricted Phenomenon? Journal of Immunology Research 2015:1–10. doi:10.1155/2015/838035

The Tabula Muris Consortium, Overall coordination, Logistical coordination, Organ collection and processing, Library preparation and sequencing, Computational data analysis, Cell type annotation, Writing group, Supplemental text writing group, Principal investigators. 2018. Single-cell transcriptomics of 20 mouse organs creates a Tabula Muris. Nature 562:367–372. doi:10.1038/s41586-018-0590-4

Tung J, Zhou X, Alberts SC, Stephens M, Gilad Y. 2015. The genetic architecture of gene expression levels in wild baboons. eLife 4:e04729. doi:10.7554/eLife.04729

Veyrieras J-B, Kudaravalli S, Kim SY, Dermitzakis ET, Gilad Y, Stephens M, Pritchard JK. 2008. High-Resolution Mapping of Expression-QTLs Yields Insight into Human Gene Regulation. PLoS Genet 4:e1000214. doi:10.1371/journal.pgen.1000214

Wang Z, Zhang J. 2011. Impact of gene expression noise on organismal fitness and the efficacy of natural selection. Proceedings of the National Academy of Sciences 108:E67–E76. doi:10.1073/pnas.1100059108

Ward MC, Gilad Y. 2019. A generally conserved response to hypoxia in iPSC-derived cardiomyocytes from humans and chimpanzees. eLife 8:e42374. doi:10.7554/eLife.42374

Westra H-J, Peters MJ, Esko T, Yaghootkar H, Schurmann C, Kettunen J, Christiansen MW, Fairfax BP, Schramm K, Powell JE, Zhernakova A, Zhernakova DV, Veldink JH, Van den Berg LH, Karjalainen J, Withoff S, Uitterlinden AG, Hofman A, Rivadeneira F, ‘t Hoen PAC, Reinmaa E, Fischer K, Nelis M, Milani L, Melzer D, Ferrucci L, Singleton AB, Hernandez DG, Nalls MA, Homuth G, Nauck M, Radke D, Völker U, Perola M, Salomaa V, Brody J, Suchy-Dicey A, Gharib SA, Enquobahrie DA, Lumley T, Montgomery GW, Makino S, Prokisch H, Herder C, Roden M, Grallert H, Meitinger T, Strauch K, Li Y, Jansen RC, Visscher PM, Knight JC, Psaty BM, Ripatti S, Teumer A, Frayling TM, Metspalu A, van Meurs JBJ, Franke L. 2013. Systematic identification of trans eQTLs as putative drivers of known disease associations. Nat Genet 45:1238–1243. doi:10.1038/ng.2756

Wright FA, Sullivan PF, Brooks AI, Zou F, Sun W, Xia K, Madar V, Jansen R, Chung W, Zhou Y-H, Abdellaoui A, Batista S, Butler C, Chen G, Chen T-H, D’Ambrosio D, Gallins P, Ha MJ, Hottenga JJ, Huang S, Kattenberg M, Kochar J, Middeldorp CM, Qu A, Shabalin A, Tischfield J, Todd L, Tzeng J-Y, van Grootheest G, Vink JM, Wang Q, Wang Wei, Wang Weibo, Willemsen G, Smit JH, de Geus EJ, Yin Z, Penninx BWJH, Boomsma DI. 2014. Heritability and genomics of gene expression in peripheral blood. Nat Genet 46:430–437. doi:10.1038/ng.2951

Yanai I, Benjamin H, Shmoish M, Chalifa-Caspi V, Shklar M, Ophir R, Bar-Even A, Horn-Saban S, Safran M, Domany E, Lancet D, Shmueli O. 2005. Genome-wide midrange transcription profiles reveal expression level relationships in human tissue specification. Bioinformatics 21:650–659. doi:10.1093/bioinformatics/bti042

Yu G, Wang L-G, Han Y, He Q-Y. 2012. clusterProfiler: an R Package for Comparing Biological Themes Among Gene Clusters. OMICS: A Journal of Integrative Biology 16:284–287. doi:10.1089/omi.2011.0118

Zerbino DR, Achuthan P, Akanni W, Amode MR, Barrell D, Bhai J, Billis K, Cummins C, Gall A, Girón CG, Gil L, Gordon L, Haggerty L, Haskell E, Hourlier T, Izuogu OG, Janacek SH, Juettemann T, To JK, Laird MR, Lavidas I, Liu Z, Loveland JE, Maurel T, McLaren W, Moore B, Mudge J, Murphy DN, Newman V, Nuhn M, Ogeh D, Ong CK, Parker A, Patricio M, Riat HS, Schuilenburg H, Sheppard D, Sparrow H, Taylor K, Thormann A, Vullo A, Walts B, Zadissa A, Frankish A, Hunt SE, Kostadima M, Langridge N, Martin FJ, Muffato M, Perry E, Ruffier M, Staines DM, Trevanion SJ, Aken BL, Cunningham F, Yates A, Flicek P. 2018. Ensembl 2018. Nucleic Acids Research 46:D754–D761. doi:10.1093/nar/gkx1098

Zhang Z, Qian W, Zhang J. 2009. Positive selection for elevated gene expression noise in yeast. Mol Syst Biol 5:299. doi:10.1038/msb.2009.58

Zhou X, Stephens M. 2012. Genome-wide efficient mixed-model analysis for association studies. Nat Genet 44:821–824. doi:10.1038/ng.2310

